# A single-mode associates global patterns of brain network structure and behavior across the human lifespan

**DOI:** 10.1101/2020.09.24.312090

**Authors:** Brent McPherson, Franco Pestilli

## Abstract

Multiple human behaviors improve early in life, peaking in young adulthood, and declining thereafter. Several properties of brain structure and function progress similarly across the lifespan. Cognitive and neuroscience research has approached aging primarily using associations between a few behaviors, brain functions, and structures. Because of this, the multivariate, global factors relating brain and behavior across the lifespan are not well understood. We investigated the global patterns of associations between 334 behavioral and clinical measures and 376 brain structural connections in 594 individuals across the lifespan. A single-axis associated changes in multiple behavioral domains and brain structural connections (r=0.5808). Individual variability within the single association axis well predicted the age of the subject (r=0.6275). Representational similarity analysis evidenced global patterns of interactions across multiple brain network systems and behavioral domains. Results show that global processes of human aging can be well captured by a multivariate data fusion approach. [147]

**Data availability:** The source data are provided by the Cambridge Aging Neuroscience Project https://camcan-archive.mrc-cbu.cam.ac.uk/. Brain data derived as part of this project and used as features for all the analyses are available on brainlife.io/pubs:

**Code availability:** Code is available on github at https://github.com/bcmcpher/cca_aging and as web services reproducing the analyes at

## Introduction

Understanding human aging and its progression in health and disease have become a critical need due to the rapidly aging world population [1,2]. Brain aging and the associated cognitive decline negatively impacts society by reducing the independence of individuals in the population, with significant costs associated with increased needs for long term support or treatment [3–7]. Costs associated with an aging population are only expected to increase over the next few years, given the increase in life expectancy [8–12]. As a result of this looming demographic shift, improving prodromal identification of individuals at risk versus individuals subject to normal aging in large human populations is becoming a priority [13–17].

The normative human aging process is accompanied by behavioral changes in performance in both cognitive and perceptual tasks across the lifespan [18–20]. Early work focussed on measuring cognitive and perceptual aging with deep characterization of a few behavioral tasks [21–24]. A staggering amount of evidence has been reported on the effect of aging to human cognition and perception. For example, language processing and letter perception deteriorate with age [25–27]. At the same time, visual contrast sensitivity, motion detection, recognition and iconic memory also decay across the lifespan [28–32]. Yet, this is not always the case as resilience to the effect of aging has been reported in a few behavioral domains such as language [26,27,33] (e.g., vocabulary size) and emotion [34,35].

Correspondingly with cognitive aging, brain aging is associated with both changes to neuronal structures and function [21,36–42]. Primary examples of changes to brain structures consist of hippocampal volume reduction [43,44], cortical thinning [45,46], and ventricular expansion [41]. At the same time, multiple examples of changes in brain functional activity have been reported [47,48]. For example, the brain hemodynamic response changes across age [49–51], prefrontal cortex activity changes during a variety of tasks [47,52,53], such as attentional control [54,55], inhibition [54,56], and executive control [19]. Finally, changes to the white matter tissue properties across the lifespan have been reported [57–64]. These studies painted a comprehensive picture of how aging affects human behavior due to alterations to either brain function and structure across multiple tissue types.

As of today, much attention has been devoted to the characterization of the changes in individual cognitive tasks and brain systems as a result of aging [21,51,65]. This approach has helped tremendously in developing an understanding of the mechanisms involved in individual cognitive functions [66–69]. Yet, a few shortcomings have been noted with single-task approaches [70,71]. For example, limits to the definition of psychological constructs [36,72], and the co-involvement of networks of brain systems in supporting individual psychological domains [62,73–75] hinder our ability to assign a one-to-one correspondence between brain systems and psychological constructs or tasks. For example, challenges in isolating cognitive processes [76–79], resulted in critiques to the very definition of psychological constructs [72]. Furthermore, studying a few constructs at a time may be also limited because multiple processes coexist within individual brain areas, as early as in the sensory systems [80,81].

Besides the mappings between a few functions and brain systems, the global patterns of change in the brain network connectivity associated with human aging, behavior, and cognition remain uncharted. To advance understanding of normative aging, we integrated multivariate [38,40,76,82–84] and network neuroscience [38,85–88] methods to develop a “many-to-many” map between behavioral domains and brain network systems. A reproducible data preprocessing pipeline was developed for the current work and made freely available as a web service on brainlife.io [89]. The pipeline was used to process data from a large sample of healthy adults (594 subjects, 18-88 years; [90]). Canonical Correlation Analysis (CCA; [83]) was then used to associate multiple behavioral measures (tasks and scales) with the connectivity properties of structural brain networks across age [73,91]. A single CCA axis of covariation successfully mapped brain networks to behavioral features. Critically, the CCA axis was associated with the individual participants’ age indicating that a coherent pattern of degradation affects both brain networks and behavior. Finally, representational similarity analysis [92] applied to the cross-validated CCA factors determined which global factors in multiple brain networks [93] and behavioral domains [90] jointly associated with predicting age.

## Results

The goal was to estimate if the structural connectivity properties of human brain networks are associated with human behavior measured from tasks, questionnaires, and scales collected across the lifespan (18-88 years, Supplemental Figure 1). To do so, data from the Cambridge center for Ageing and Neuroscience was used, the dataset is hereafter referred to as CAN [90]. The CAN dataset contains a deeply phenotyped cohort of cognitively healthy individuals (594 used here; See **Methods** for inclusion criteria) evenly sampled across the lifespan (100 subjects per decade). We used 388 behavioral scores from 33 assessments published with the CAN dataset. These scores consisted of either reaction times, accuracy, or performance scales–see example histograms of each type of score plotted across age are shown in **Figure 1a** and **Supplemental Figure 1**. A total of 334 normalized behavioral measures were extracted and utilized for all subsequent analyses (140 reaction-time, 188 accuracy, and 6 assessments; see **Methods: Performing the CCA**). The CAN behavioral measures were originally mapped into five behavioral domains [90]: attention, language, memory, motor, and emotion. In addition to these, scores from social and clinical assessments were organized into two additional domains for a total of 7 behavioral domains (**Figure 1a**).

**Figure 1.**
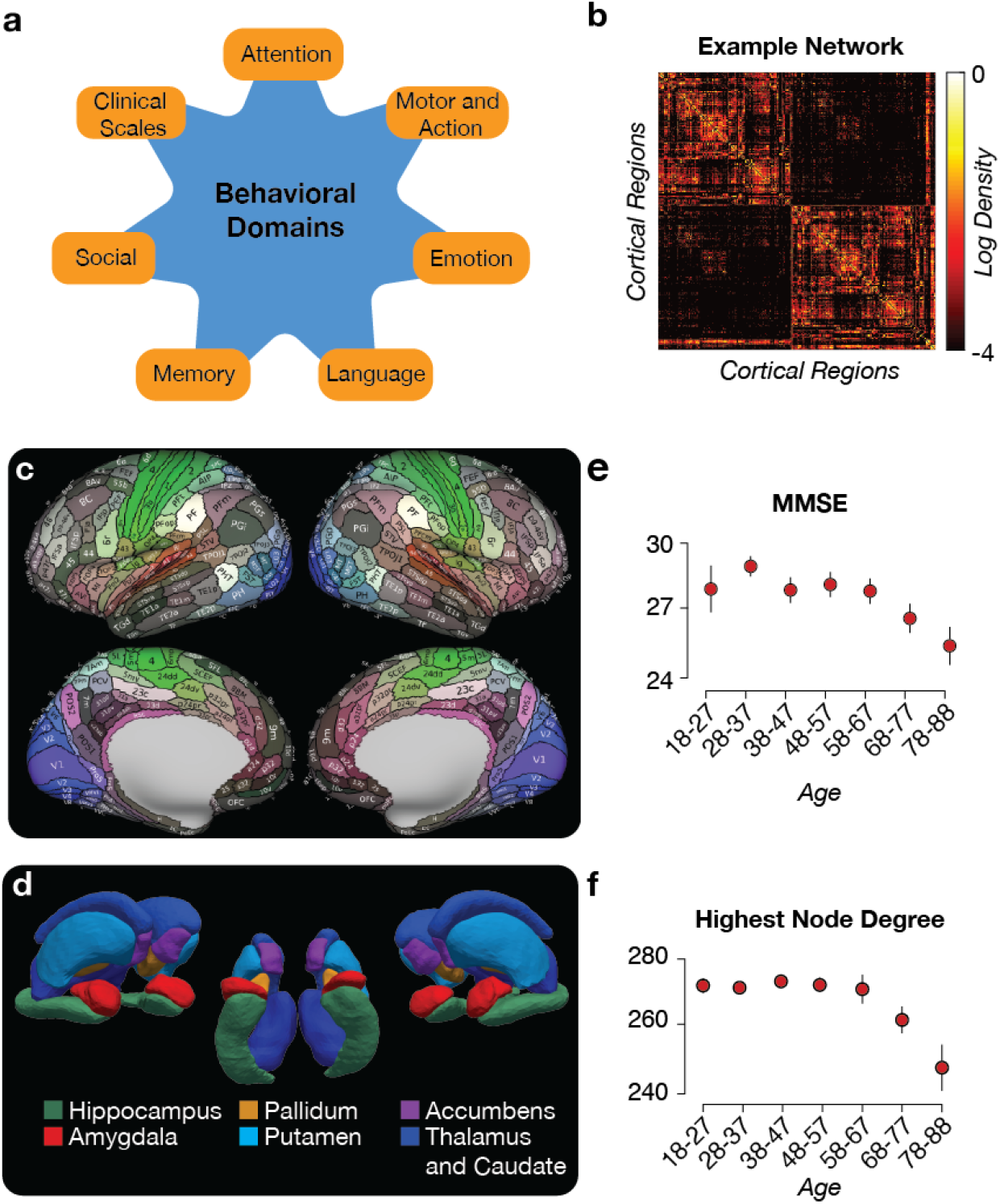
Main Findings. **a. The behavioral domains.** A simple graphic displaying the different behavioral domain labels is displayed. **b. An example network.** This is an example of an individual’s network that is created for analysis. Each row and column represents a cortical region while the color represents the log scale of the density of the connection between nodes (weaker connections are black, stronger connections are yellow). **c. The cortical regions used to create the network structures.** This figure, taken from Glasser et al. (2016), represents the cortical labels used to construct the network in **b. d. Additional subcortical labels.** A surface rendering of the subcortical labels that were estimated and added to the cortical labels shown in **e. Trends in behavior over the lifespan.** The mean and standard deviation for the Mini-Mental State Exam (MMSE) is binned into decades and is displayed. Error bars represent 2 units of standard error. **f. Trends in the connectome over the lifespan.** The mean and standard deviation for the highest node degree is binned into decades and is displayed. Error bars represent 2 units of standard error. Subjects were binned by decades starting at 18 years of Age (see [90]).

Whole-brain structural networks were estimated using diffusion-weighted magnetic resonance imaging (dMRI) data from 594 subjects [90]. Neuroimaging data were processed using an automated and reproducible pipeline using brainlife.io (see **Table 1**; [89]). **Figure 1** panels **b-d** show a representative network, as well as the cortical (MMP v1.0; [94]) and subcortical [95] atlases used for network neuroscience data generation. A total of 376 regions (366 cortical and 10 subcortical) were used to generate each network. The dMRI data was processed for artifact removal and fiber model fitting using a recent robust method (see [96] and <u>bl.app.68</u>). Whole-brain tractograms were generated using a novel pipeline called Reproducible Anatomically Constrained Ensemble Tracking (RACE-Track; see <u>bl.app.101</u>), which integrates two established methods [97,98]. RACE-Track tractograms were combined with the regions of interest from the two atlases (see <u>bl.app.23</u>) to build individual subjects’ connectivity matrices. Connections present in less than half of the subjects were eliminated (see also **Supplemental Figure 2a**; [99]). Connection density (*C_d_*, see *Eq. 1,* **Figure 1b,** generated by web service <u>bl.app.121</u>; [100]) was used as network edge weight. The 70,500 *C_d_* estimates were reduced to 376 node degree estimates used for all subsequent analyses (see **Supplemental Figure 2a** [91]).

**Table 1.** Open brainlife.io web services and containerized applications implementing the processing pipeline developed for the current work. We developed a computationally reproducible data processing pipeline utilizing the cloud computing platform brainlife.io [89]. Each step within the pipeline is described in **Table 1** and available for download as Docker [137] container run via Singularity [138]. Each step of the pipeline is also publicly shared as an App (web service) on the brainlife.io platform. brainlife.io Apps can be freely executed on public or private computing resources. All processed brain imaging data and Apps are accessible in a single record as a brainlife.io publication at <<DOI-TO-MY-DATA>> [89].

| App name | App purpose | DOI |
| --- | --- | --- |
| AC-PC Alignment of T1 | *Align T1w anatomical images along the AC-PC plane* | bl.app.99 |
| AC-PC Alignment of T2 | *Align T2w anatomical images along the AC-PC plane* | bl.app.116 |
| FreeSurfer (v6.0.0) | *Create surfaces and cortical labels* | bl.app.0 |
| dMRI Preprocessing | *Correct dMRI data of artifacts and align to T1w image* | bl.app.68 |
| RACE-Track (Tractography) | *Create an anatomically constrained whole brain ensemble tractogram* | bl.app.101 |
| Multi-Atlas Transfer Tool (maTT) | *Map MMP atlas to individual subjects' brains* | bl.app.23 |
| Network Construction | *Create networks from tractography and cortical labels* | bl.app.121 |

## Changes in network neuroscience and behavioral measures across the lifespan

The relationship between networks’ node degree and performance in behavioral tasks and assessments was explored across the lifespan (**Figure 1e**, **f** and **Supplemental Figure 1**). The goal was to ensure replication of known trends established in the literature. Changes in both network and behavioral measures were observed – especially starting at the fourth-decade **Figure 1e, f** [57].

A large subset of behavioral measures varied as a function of age (**Figure 1** and **Supplemental Figure 1**). Reaction times increased with age, accuracy decreased, and performance in multiple scales either decreased or increased with age depending on the scale valence ([101,102]; **Supplemental Figure 1a**). For example, reaction-time increased in the Force Matching Task, but accuracy decreased (**Supplemental Figure 1a**), performance in the Mini-Mental State Exam (MMSE) also decreased (**Supplemental Figure 1a**; [90]). To approximate these trends we fit a quadratic polynomial to each behavioral measure. We found that 166 of the total 334 behavioral variables increased with age (quadratic term: 0.034 ± 0.083 s.d., R^2^ = 0.957 ± 0.123 s.d., AICc = 49.067 ± 4.977 s.d.), and the remaining 168 decreased with age (quadratic term: −0.005 ± 0.016 s.d., R^2^ = 0.988 ± 0.050 s.d., AICc = 48.232 ± 1.079 s.d.).

To evaluate whether the brain networks generated using RACE-Track showed sensible changes in properties across the lifespan, we estimated the highest node degree (**Figure 1f** and we observed similar patterns when evaluating network density and efficiency see **Supplemental Figure 1b**). To summarize the effect of age on these three measures, we fit a quadratic polynomial. The highest network node degree was well described by a negative quadratic term; it increased early and decreased later in life (quadratic term: −0.005 ± 0.008 s.d., R^2^ = 0.9996 ± 0.0004 s.d., AICc = 48.007 ± 0.007 s.d.; network density and efficiency demonstrated a similar negative quadratic pattern, **Supplemental Fig 1b**, *right*).

## A single-mode of covariation relates individual differences in structural networks and behavior with subjects’ age

After replicating the established quadratic trends in brain and network properties and behavioral variables with age, we set out to build a model that would linearly associate all the measurements from the two data domains – brain and behavior. We did this because we wanted to determine whether a linear association well approximated the relationship between brain networks and behaviors. To do so we used canonical correlation analysis (CCA; [83]). CCA finds the linear combination of variables that best associates measures from the two data domains across subjects. In the following analyses, CCA found the best linear combination of 376 brain network properties and 334 behavioral measures (see **Figure 2**, **Methods**, and **Supplemental Figure 2a**, **b** and **c**). To do so, the behavioral measures and networks’ node degree estimated in each subject were organized into two matrices (*D_1_* and *D_2_*; **Supplementary Figure 2a**). Confounding variables such as sex, handedness, height, body weight, heart rate, and blood pressure were regressed out from both D_1_ and D_2_ (see **Supplementary Figure 2a**, **Methods**, and [83]). The eigenvector matrices (*E_1_* and *E_2_*) estimated from *D_1_* and *D_2_* via Principal Component Analysis (PCA) were used as inputs to CCA ([83]; **Supplemental Figure 2b** and **c**). CCA found the weights (*a* and *b*) and canonical factors (*F_1_* and *F_2_*) that best approximated *E_1_* and *E_2_* (**Supplemental Figure 2c** and **d**).

**Figure 2.**
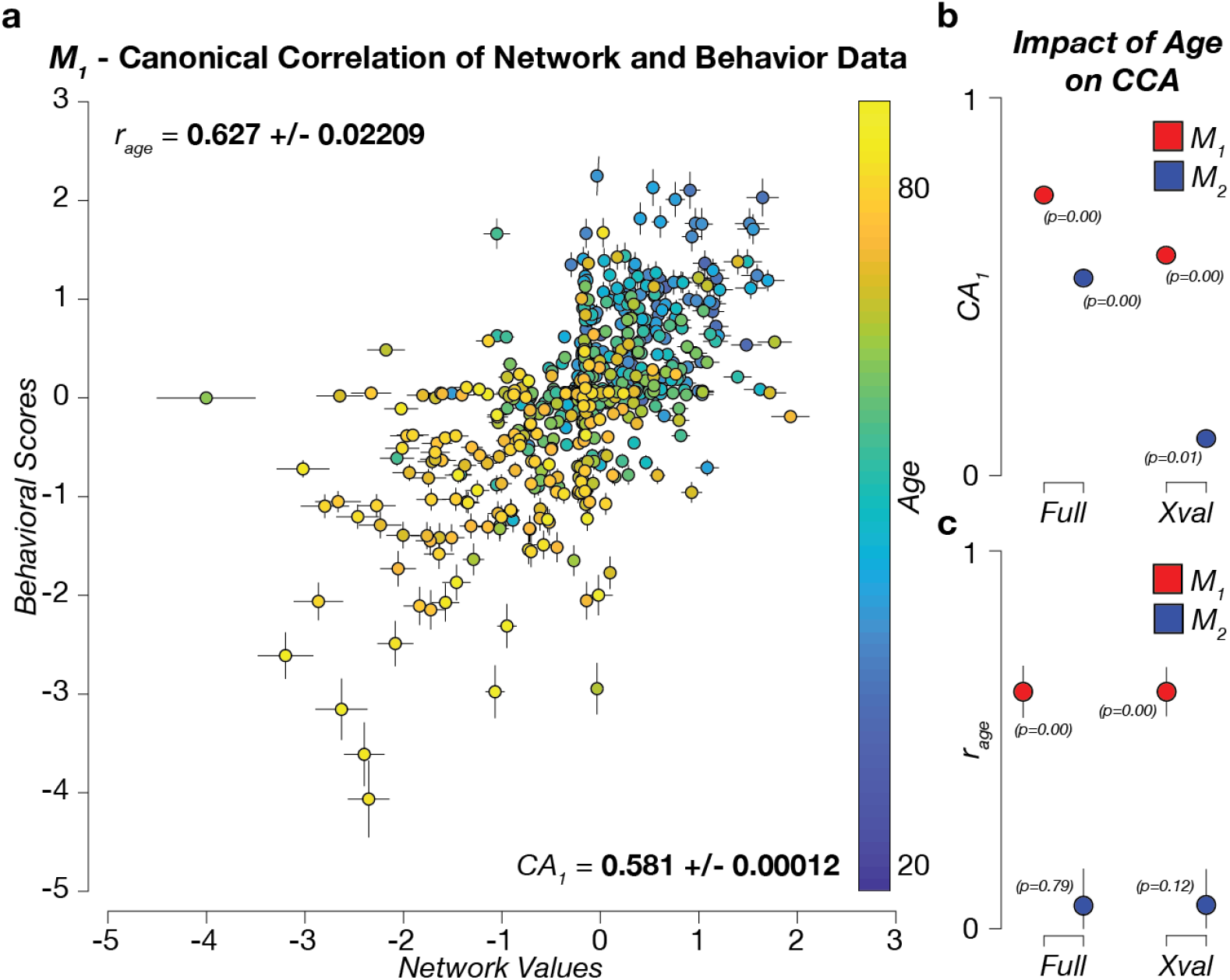
Human age explains the majority of the variability in the linear association between brain networks and behavioral variables. **a.** The first canonical axis (CA_1_) from the cross-validated model (*M*_1_). Each point represents a subject's cross-validated factor score on the brain and behavior axes, respectively. Error bars represent ±3 Standard Error of the Mean (SEM). The color of each point represents the age of the subject. The labeled correlations refer to the correlation of age to the combined canonical axis (*r_age_*; top) and the correlation between the two canonical axes (*CA_1_*; bottom) ±3 SEM. **b.** Association between the brain network and behavior data for the first canonical axis (CA_1_) shown for both *M*_1_ and *M*_2_. Note the reported ±3 SEM are smaller than the symbols. **c.** Association (*r_age_*) between the data in the first canonical axis (CA_1_) and each participant’s age. Error bars show ±3 SEM. Statistical significance estimated by bootstrap.

A first CCA model (*M*_0_) used a large number of PCA eigenvectors (100) as inputs and no cross-validation to evaluate the best possible fitting model to the data [83]. *M*_0_ returned multiple large and statistically significant modes of covariation explaining the relationship on the first canonical axis (CA_1_) between networks and behaviors (CA_1_=0.860-0.1072; *p_1-75_*=0.000 and *p_76-85_*<0.01 bootstrap test). After this exploratory model, a 5-fold cross-validation approach (subjects as a random variable; **Supplementary Figure 2e**) was used to further test the association between networks and behavioral variables. A grid-search approach was used to generate 9,801 CCA models with different combinations of PCA numbers (spanning from 2 to 100). Each model was cross-validated using 15,000 5-fold throws for a total of 147,015,000 tested models. The first canonical correlation for a majority of these models was large with a mean CA_1_ of 0.5468±0.09 s.d. (min and max 0.235 and 0.604, respectively). Over 90% of the CA_1_ values across all models lay above 0.55, this indicated that, with a few exceptions, a majority of the cross-validated models CA_1_ explained the data reasonably well.

Because of the established association between age and several brain and behavioral-measures (see **Figure 1**), we selected the model (*M*_1_) with the highest correlation between age and CA_1_ (*r_age_*=0.627±0.02209). *r*_age_ was computed using a multiway correlation via a general linear model where subjects’ age was predicted using CA_1,behavior_, and CA_1,network_ as regressors (see **Methods**, *Eq. 3*). The selected model, *M*_1_, had 38 brain and 40 behavior principal components and a significant CA_1_ of 0.581±0.0001 (*p*=0.000 bootstrap test). Noticeably, CA_1_ was the only significant mode in *M*_1_ (see **Figure 2a** and **b**, see also **Methods** and **Supplemental Figure 2f-h**). The remaining modes were either not statistically significant (average CA_3-38_=0.004±0.014; *p*=0.442±0.254 s.d. bootstrap test) or had no statistically significant loadings (CA_2_; see **Supplemental Figure 2i** and **j** and next section). *r*_age_ was strong and statistically significant for the first mode as expected given the model selection procedure (0.627±0.022; *p*=0.000 bootstrap test) and not significant for the rest of the modes (average *r*_age_ for CA_2-38_ was 0.077±0.022; *p*=0.46±0.029 bootstrap test).

Two additional analyses we performed to further explore and validate the results. First, *r*_age_ was also computed for the exploratory mode, *M*_0_. In this case the association was found to be significant for CA_1_ only (*r*_age_=0.611±0.023; *p*=0.000 bootstrap test; average CA_2-38_, *r*_age_=0.063±0.015; *p*=0.726±0.265 s.d. bootstrap test, see **Methods**). Finally, to validate the consistency of our approach, and the role of age in the association between brain networks and behaviors a new CCA model was specified (*M*_2_). *M*_2_ was similar to *M*_1_ with the exception that the subjects’ age was regressed out from both the brain and behavior data before performing the PCA (See **Methods**, *Eq. 2*). CA_1_ for *M*_2_ was extremely small even though somehow significant (0.098±0.0016; *p*=0.007 bootstrap test). None of the remaining modes for *M*_2_ were significant (average CA_2-38_, 0.0003±0.017; *p*=0.520±0.290 s.d. bootstrap test; see **Supplemental Figure 2k, l**). As expected, the correlation between age and CA_1-38_ for *M*_2_ was insignificant (average *r*_age_=0.075±0.019 s.d. and *p*=0.490±0.300 s.d. bootstrap test). These results show that the quadratic trends for individual variables across the lifespan (as shown in **Figure 1e, f**) in the two data domains (behavior and brain networks) are well coupled by CCA into a single linear trend. In addition, the CCA mode is also strongly associated with the participants’ age.

To summarize this main result, we implemented a hypothesis-driven approach to CCA using cross-validation and tested our hypothesis of an association between age and the CCA model. More specifically, we tested the degree to which the participants’ age predicted (in cross-validation terms) the variability in the linear combination between hundreds of variables from brain and behavioral measures. The results show that a major portion of the variability in the CCA model is effectively associated with age. This result does not mean that other variables were not associated with the model, indeed they were, as the CCA model is significant and correlation in CA_1_ is strong.

## Brain network and behavioral components contributing to the CCA

We further investigated the behavioral domains and brain network nodes that most contributed to *M_1_*, and indirectly to the predominant association with the subject’s age. To do so, the CCA variable loadings, *L*_1_ and *L*_2_ were estimated by computing the correlation between each column in *D*_1_ and *D*_2_ and every column in *F*_1_ and *F*_2_, respectively (**Supplementary Figure 3a**; [83]).

The top 20 *L*_1_ and *L*_2_ are reported for CA_1_ (**Figure 3a** and **b** for brain networks and behavior respectively). The results show that the brain network nodes known to be affected by aging were among the top contributors [34,65]. The inset in **Figure 3a** shows the mapping of all the CA_1_ loadings to the cortical and subcortical surfaces. The visualization of the loadings shows that the distribution of loadings is homogeneously distributed with a few hot-spots, with positive foci in the frontal lobe, the hippocampus, the putamen and portions of the default mode network. In the next paragraph, we summarize how the brain area loadings relate to previously reported functional network labels [93]. In parallel, behavioral tasks and scales measuring cognitive and emotional domains also known to be affected during human aging returned meaningful loadings as well [18,66]. These domains consisted of visual recognition, attention, and memory tasks, as well as, reasoning and language comprehension scales. No significant canonical loadings were found for CA_2-38_ (see **Supplemental Figure 3b** for *CA*_2_ loadings). For completeness, we also tested all the loadings for cross-validated *M*_2_. No significant loading was found for this model (see **Supplemental Figure 3c** p>0.05 bootstrap test). In sum, none of the CA in *M*_2_ was interpretable, hence the model was uninterpretable.

**Figure 3.**
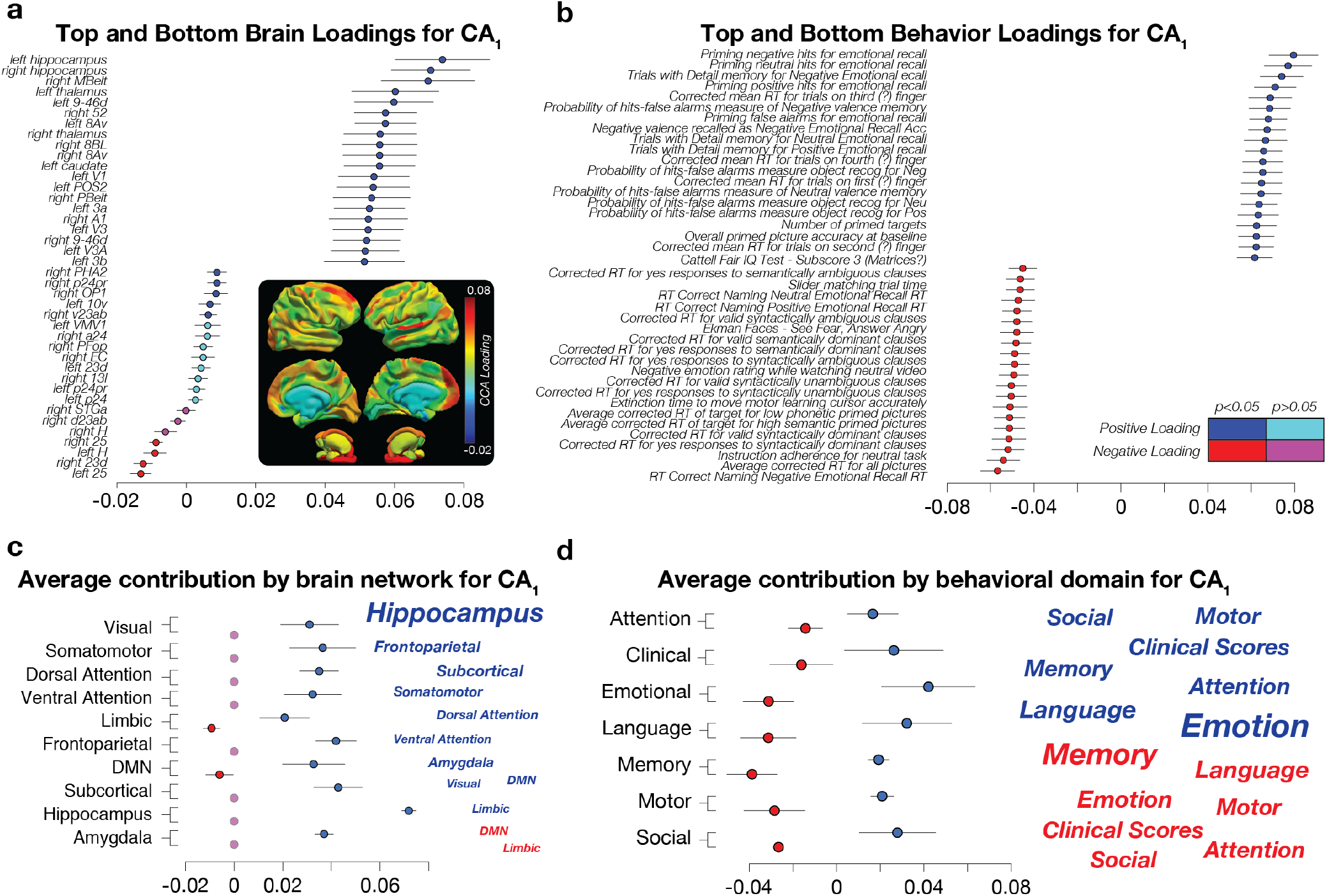
Top brain network nodes and behavioral variables contributing to the canonical correlation analysis. **a**. **The highest and lowest contributing cortical nodes within the network**. The top and bottom 20 CA_1_ brain regions sorted by loading magnitude. Positive and significant loadings are shown in blue, negative in red. Non-significant loadings are shown in cyan or magenta. The anatomical inset visualizes the CA_1_ loadings projected on the cortical and subcortical regions used to build the networks. **b. The highest and lowest contributing behavioral scores within the domain**. The top and bottom 20 CA_1_ behavioral scores sorted by loading magnitude (same conventions as in **a). c. Average CA_1_ loading by the major functional brain network as described by Y_2011_ [93]**. The word cloud in the right-hand panel shows the functional network name scaled by contribution to CA_1_ (the larger the font the greater the contribution). **d. Average CA_1_ loading by behavioral domain**. The word cloud in the right-hand panel shows the domain scaled by contribution to CA_1_. All symbols report mean±2 standard errors (s.e.) estimated across 10,000 cross-validation throws of *M*_1_.

To summarize the top contributors to *M_1_*, word-cloud representations of the variable domains were created (**Figure 3c** and **d**). The word-cloud for the brain network loadings was built by assigning the labels from the Glasser atlas to a set of established functional network labels [93], hereafter referred to as Y_2011_. Each of the 376 nodes in our networks was assigned to one of the seven functional networks in Y_2011_; Visual, Somatomotor, Limbic, Ventral attention, Dorsal Attention, Frontoparietal, Default Mode (DMN; see **Methods** for the assignment of the network’s nodes to major functional networks in Y_2011_). In addition, the hippocampus, amygdala were kept separated and all the remaining subcortical structures were combined and reported as subcortical (i.e., pallidum, putamen, accumbens, thalamus, and caudate). This process generated a total of ten words (or clusters). Each word was scaled by the sum of the loadings of all the nodes assigned to the cluster (**Figure 3c,** see also **Methods** for a description of how the word-clouds were generated from the loadings).

This analysis highlighted the top functional networks and subcortical structures contributing to *M_1_*, the one that indirectly also contributed to predicting the subjects’ age. The limbic and default-mode networks and the hippocampus returned the strongest contribution (positive and negative, respectively). Other networks, such as the frontoparietal and dorsal attention, as well as the amygdala, provided a strong positive contribution. A similar, word-cloud representation was implemented also for the behavioral variables. Each of the 334 tasks and scales was uniquely assigned to one of the seven behavioral domains described in **Figure 1a.** Their loadings were then averaged across all tasks and scales within each domain (**Figure 3d**). The word-cloud summary representation for the behavioral loadings shows the variables in the emotional, language, and memory domains contributed the most to *M_1_*. Other variables such as social, attention, motor, and clinical scores provided secondary contributions. Next we evaluated whether the properties of the rich club properties of the brain networks [73,103,104] were associated with the CCA loadings.

## The brain rich-club contribution and canonical correlation

One of the most reliable findings in network neuroscience is the brain rich-club organization [73,103,104]. Here, we were interested in estimating whether there was a relation between the top contributing brain regions in CA_1_ and the regions participating in the rich-club organization of the brain. More specifically, we evaluated the extent to which the top CA_1_ brain loadings (i.e., **Figure 3a**) mapped onto the rich-club core or periphery. To do so, the mean network across subjects, *N*_μ_, was generated by averaging streamline density for each edge (edges not appearing in at least 50% of the subjects were set to 0 [91]). The number of nodes’ participating in the rich club depends on the brain parcellation used. In our study, the rich club core was defined as the top 15% highest-degree nodes in *N*_μ_. This proportion of rich-club nodes was previously reported by [103] (see also **Methods** for more details). Fifty-four regions from the HCP-MMP (v1.0) parcellation and subcortical labels were assigned to the rich-club core. The remaining 376 regions were assigned to the rich-club periphery (**Figure 4a** *blue* and *gray* respectively). We found that this procedure successfully assigned regions to the rich club nominally matched the regions reported in the literature (e.g., superior parietal, precuneus, superior frontal cortex, putamen, hippocampus, and thalamus; **Figure 4a**; see also [103] for a comparison).

**Figure 4.**
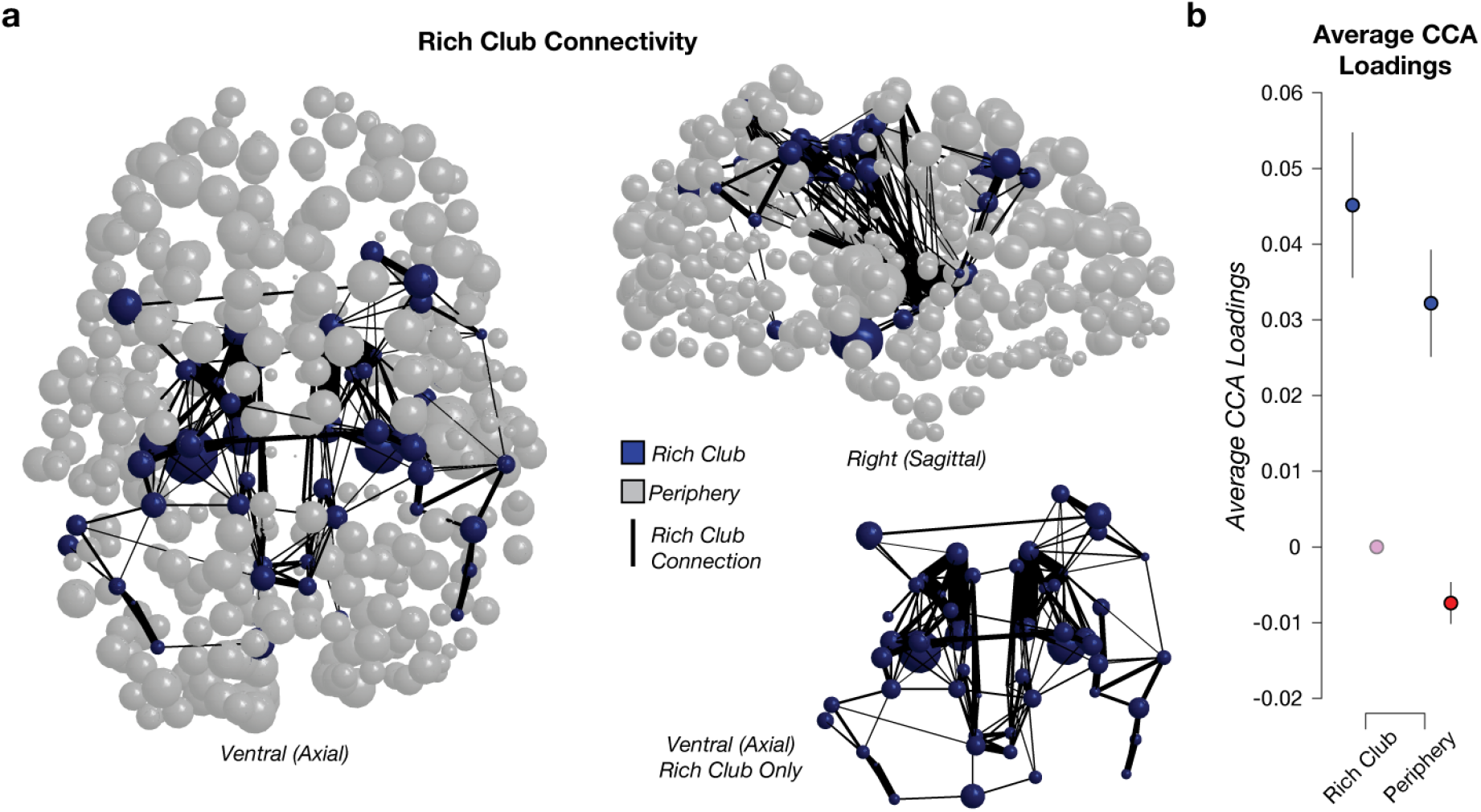
Relationship between the brain connectome rich-club and the contribution to the CCA. **a. A ball-stick representation of the rich club scaled by CCA loadings**. Axial and sagittal views of the relation between rich club and CCA loadings. Each brain region is displayed as a sphere. Blue spheres are part of the core rich-club. Gray spheres are part of the periphery of the rich-club. The diameter of each sphere was scaled by cross-validated CA_1_ loading in M_1_. The black lines show connections between rich club nodes. The thicker the line, the higher the average streamline density. The right-hand bottom ball-stick representation shows an axial view of the rich club nodes and connections isolated from the periphery. **b. The average CA_1_ loading across rich club core and periphery**. Positive and negative CA_1_ loadings for rich club core and the periphery (mean ±2 s.e. via cross-validation of model M_1_).

After defining the rich-club core and periphery, we correlated each node’s loading on CA_1_ with the normalized node degree used to estimate the rich club participation. The CA_1_ loadings for regions within the rich-club core or periphery were averaged together. On average, the loadings were higher for the core than the periphery (0.045±0.0096 s.e., *p*=0.00 and 0.032±0.0071 s.e., *p*=0.00, respectively). This shows a trend for higher CA_1_ loadings within the rich club, even though no significant difference was found in the mean loadings between core and periphery (**Figure 4b**; *p =*0.722 bootstrap test). Supplemental **Figure 4** shows the rich club participation coefficient and CA_1_ loadings for each brain region color-coded by their participation to the rich club or periphery.

We also tested whether the measure used to define the rich club (the node degree of the average network) was significantly correlated with the brain variable loadings in the network axis of CA_1_ (for reference the values shown in **Figure 3a**). We report two findings. First, a medium-strong correlation was found if the rich-club organization was disregarded and the correlation computed across all brain regions (r=0.642±0.034 s.e., Spearman rank r; *p =* 0.00 bootstrap test). The correlation was weaker when we considered only the 54 regions within the rich-club core (r=0.26±0.127 s.e., *p*=0.0283), and larger when we considered the 322 regions in the rich-club periphery (r=0.56±0.041 s.e., *p*=0.00). Overall, the results show an interesting trend with higher CA_1_ loadings for regions within the rich club compared to regions in the rich-club periphery (a statistically non-significant trend) and a statistically significant correlation between the node degree in the rich-club periphery. The results can be interpreted as indicating a heterogeneous contribution of regions within and outside the rich-club in the association between brain and behavior and in predicting age.

## Describing the multivariate relationships between networks and behaviors using representational similarity analysis

Our overarching goal was to describe a multivariate fingerprint of the associations between brain networks and behavioral domains. Most previous analyses looked at pairs of associations to characterize the changes in individual cognitive tasks and brain systems as a result of aging [21,51,65]. This approach focussed on understanding the relationships between individual cognitive functions and brain systems across the lifespan [66–69]. As a result a global fingerprint of the multivariate changes in the brain network connectivity, behavior, and cognition associated with human aging, have not been described. We used Representational Similarity Analysis (RSA; [92,105]), to develop a “many-to-many” map between behavioral domains and brain network systems.

Our analyses in the previous sections focussed on *M*_1_ to establish the relationships between brain and behavior across the lifespan and how fundamental brain network properties contribute to such relationships. Hereafter, we focussed on the multivariate relationships between brain network and behavior domains. To do so, we derived an approach that used the CCA variables loadings (*L_1_* and *L_2_*, **Supplemental Figure 5a**) as inputs to RSA. An RSA typically quantifies the dissimilarity between variables from two data domains; for example and more commonly, brain function and behavioral performance [92,105]. In our application instead of using the data from the two domains directly, we used the variable loadings estimated for each canonical axis in *M*_1_. More specifically, every variable loading in *M*_1_ (376 network and 334 behavior variables) was correlated with the loadings of all the remaining variables, and dissimilarity was then computed using *Eq. 4*. This process generated a square, symmetric RSA matrix, *S*_1_, of size 710×710.

This approach leverages the inherent structure of the data captured by the CCA model to describe how the loadings of each brain and behavior variable are associated amongst themselves. The assumptions of this analysis are that the CCA variables loadings contain a fingerprint of the multivariate associations of the brain and behavioral variables. The approach allowed us to model (1) the brain network-to-network similarity, (2) the behavior-to-behavior similarity, as well as (3) the brain network-to-behavior similarity. *S*_1_ was summarized by averaging the RSA values from the individual brain network nodes and behavioral variables within the 10 brain networks of Y_2011_ and 7 behavioral domains (as described in the previous section and in **Methods**). This reduced the dimensionality of the symmetric dissimilarity matrix from 251,695 (upper diagonal of the original 710×710 matrix) to 136 (upper diagonal of the 17×17 matrix; see **Supplementary Figure 5c**). *S*_1_ can be divided into three primary regions; brain network-to-network (*ntn*), behavior-to-behavior (*btb*), and behavior-to-network (*btn*). Because the interest here was to capture the unique relationship between brain and behavioral variables, *btn* was the focus of all the following analyses (**Supplementary Figure 5d**). The results were visualized using a modified chord plot that allowed us to show how multiple associations between brain and behavior load onto *M*_1_ simultaneously (**Supplementary Figure 5e**; see **Methods**). The chord plot was generated by thresholding the RSA values in *btn* the top quartile. Two aspects of the chord plots shown in **Supplemental Figure 5e** are of interest. First, the number of domains (or networks) that each network (or domain) contributed to is described by the size of the peripheral segments for each network and domain. The larger the size of the segments, the more contributions of a network (domain) to the various other domains (networks). Second, the associations between each functional network and behavioral domains are described by individual chords (if a chord exists two domains or networks interacted in the CCA).

## The difference between RSAs shows multivariate brain-behavior associations the variability accounted for by age

Hereafter, we wanted to further emphasize the contribution of the individuals’ age to the RSA. To do so, we used the second CCA model, M_2_, in which participants’ age was subtracted out (deconfunded). An RSA model was set up using the variable loadings in M_2_ to create S_2_. The impact of age was then further isolated by subtracting S_2_ from S_1_ to compute S_d_ (**Figure 5**). S_d_ describes the multivariate (many-to-many) associations between the networks and behavioral domains after the variables associations independent of age have been removed. In other words, the matrix, *S*_d_, evidences the dissimilarity values that are related to age but not to the rest of the variables in the model because any common dissimilarity was subtracted out. Because the interest was to capture the unique relationship between brain and behavioral variables, the following description focuses on the cross data-domain interactions (*btn*; **Figure 5b**). A chord plot was used to describe the many-to-many relationships (**Figure 5c**).

**Figure 5.**
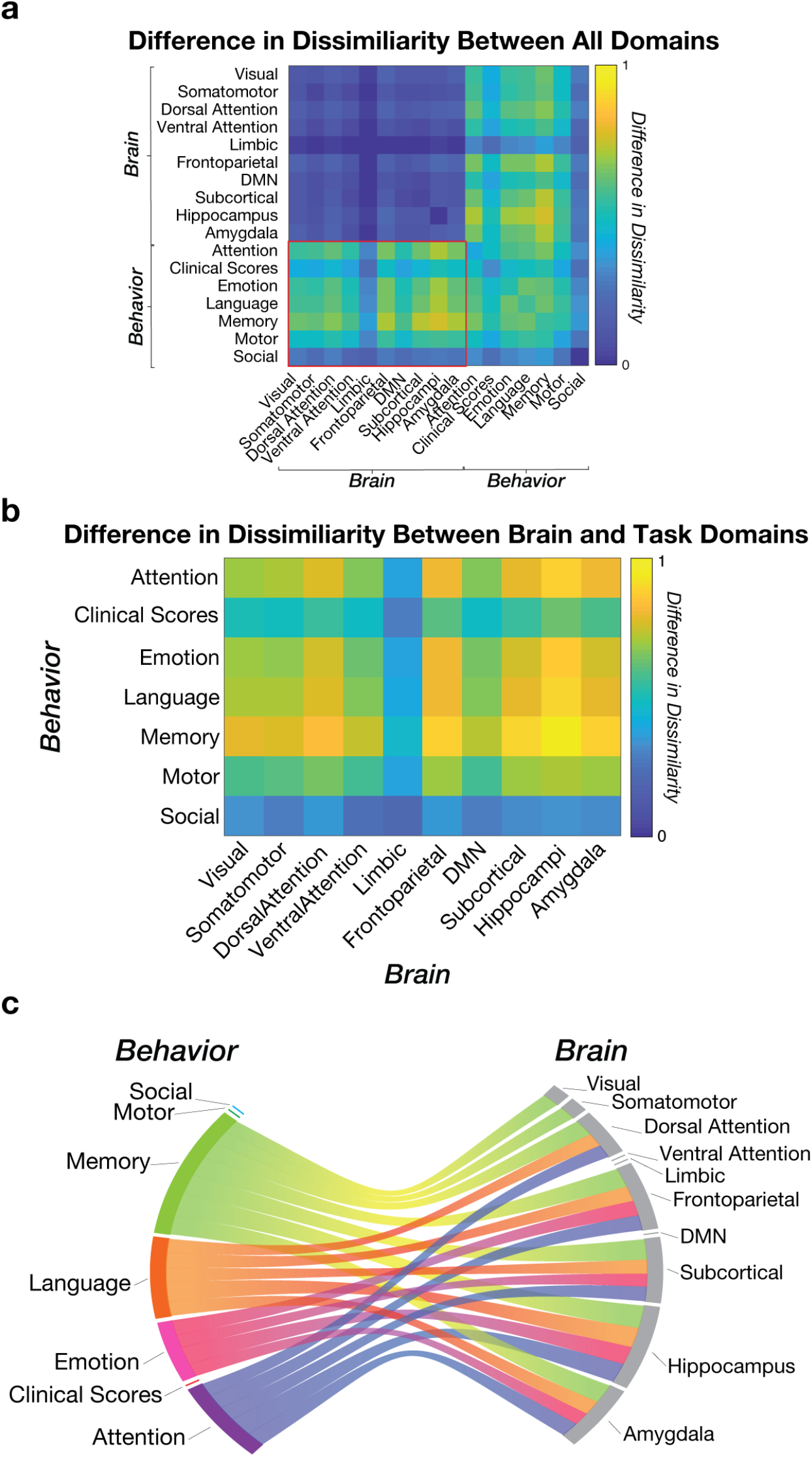
Difference in Representational Similarity Analysis (RSA) shows many-to-many associations between the modules from the brain networks and behavioral domains. **Differences in RSA (S_d_) between two CCA models (M_1_ and M_2_).** The values in the matrix S_d_ represent the difference between S_1_ and S_2_. The higher the RSA value, the higher the impact of age. The red rectangle represents the subset of brain-behavior interactions emphasized in **b** (referred to as *btn* in the main text). **b. Subset of the multivariate interactions between brain networks and behavioral domains.** The data in panel **b** shows a subset in panel **a** emphasized by the red rectangle, also referred to as *btn* in the main text. This subset shows the multimodal dissimilarity between brain and behavior domains across the lifespan. **c. Top quartile of the differences in dissimilarity between brain networks and behavioral domains.** This modified chord plot represents the top interactions between the brain networks and behavioral domains in *btn*. Associations below the 75th percentile were removed from the visualization. Two aspects of the chord plots can be appreciated. First, the number of domains or networks that each network or domain contributes to is shown by the size of the peripheral wedged-segments for each network and domain. The larger the size of the segments, the more contribution of a network (domain) to the various other domains (networks). Second, the associations between each functional network and behavioral domains are described by individual chords (if a chord exists two domains or networks interacted in the CCA).

The results of this RSA analysis were consistent with previous reports [34,38–40]. More specifically, results showed that the behavioral domains most affected across the lifespan were memory, language, emotion, and attention (i.e., the behavioral domains passing the top 25% cut-off threshold; **Figure 5c**). Going beyond previous reports, the RSA results showed that the top behavioral domains were associated not just with a specific brain network, but with multiple of them. More specifically, memory concurrently associated through the lifespan with the hippocampus (top association), the subcortical structures, the frontoparietal network, amygdala, dorsal attention network, the visual, somatomotor networks but the ventral attention, default mode and limbic networks (see **Supplemental Table 1**). Emotion was associated through the lifespan with changes in the subcortical structures but also by the frontoparietal network, the hippocampus, and the amygdala. Similarly, attention was affected by the dorsal attention and frontoparietal network but also by the amygdala, hippocampus, and subcortical structures. Language and memory show a similar pattern, with moderately high associations between many brain networks with notably strong connections to the hippocampus. Whereas some of the previous work has been devoted to characterizing individual cognitive tasks and brain systems in relation to aging [21,51,65], the current analysis showed that multiple brain networks meaningfully contribute to the variation in behavioral performance and cognition across the lifespan. The many-to-many relationships between brain networks and behaviors reported here are unexplored but critical to understanding the process of aging in normal and diseased populations.

## Discussion

Brain aging and disease have profound effects on society. Because of the increasing expected lifespan of the world population –many of us are living longer– it is increasingly important to understand how we can age healthily. Costs associated with an aging population are only expected to increase over the next few years, given the increase in life expectancy (1–5). In Europe, estimates reach up to 200 million individuals affected yearly, directly or via caregivers’ networks. Such impact accounts for up to 800 billion euros in annual costs. For example, brain disease is likely to affect up to 25% of the European population and nearly 38% of the remaining global population. These numbers make understanding the brain and its relation to behavior paramount to the well being of society across the globe. Critical to this goal, is to increase our understanding of the normal variability across individuals in the large population is fundamental to improve the prodromal identification of individuals at risk versus individuals subject to normal aging (6–10). To address the need to capture human variability and aging at least three aspects of scientific inquiry will need to be addressed. First, more high-quality data will need to be connected to reach a population-level understanding of the human brain and behavior. Second, advanced computational approaches will need to be developed that can actually exploit the value of the data at the right scale. Finally, infrastructure will need to be in place to support large-scale computational analysis of thousands of individuals.

To understand aging we need large-scale, population-level studies. There is an increasing need to better understand the trajectory of healthy aging as a large portion of the world’s population rapidly progresses into old age. The need to understand not only the expected course of senescence, but the interaction of brain structure and behavior will become increasingly important to understand the myriad of conditions that can impede independence later in life. Early open projects of this nature, such as the Alzheimer’s Disease Neuroimaging Initiative (ADNI), have required large publicly available tools to facilitate the larger number of researchers who have analyzed factors related to ones in this dataset [15,16,106–109]. More recent, large-scale population projects such as the Human Connectome Project, the ENIGMA Consortium, and the UK Biobank [60,110,111] have collected, organized, and openly distributed data to allow the implementation of data-driven neuroscience methods, mine novel findings, and build normative models of brain structure and function [94]. These projects are already pursuing similar methods, and the presented analysis could easily be applied and extended to these datasets [38,60,71,83,110].

To fully exploit large scale datasets advances in both computational methods and data-analysis infrastructure are needed. We described a multivariate analysis of a multimodal dataset [90]. Multiple measures of brain structural connectivity and behavior were combined to estimate the impact of human age on multiple brain systems and behavioral domains. A model testing framework was developed using cross-validation to demonstrate a strong linear association between brain connectivity and behaviors. The majority of the variance in the linear association between connectivity and behavior was accounted for by the first mode of the model. This model was selected to maximize the correlation with subject age. Data derivatives (https://doi.org/10.25663/brainlife.pub.21), code (https://github.com/bcmcpher/cca_aging) and reproducible cloud services (See **Table 1**) developed for the present work are made available for reuse by the wider scientific community. Finally, the RSA analysis showed a series of global patterns of association between the model, selected to maximize the correlation with the subjects’ age, and that resemble findings reported when individual variables are measured individually.

In previous studies, most commonly, a few behavioral or psychological variables have been singled out and related to the changes in brain networks to capture the aging process across the lifespan [21,51,65]. Describing how many of these cognitive and behavioral variables change across the lifespan has been a challenge [62,73–79]. For example, memory has been related to hippocampal volume and reaction time [43,44,78]. However, these targeted studies run the risk to miss that the hippocampus also has significant contributions to emotion processing and guiding attention [28,34]. Here, we proposed an approach that allowed us to represent the changes in multiple brain network systems and behavioral domains simultaneously. Our results provided a new way of quantifying the multivariate relationships to understand how these variables simultaneously change across the lifespan. The results show the changes in behavioral and psychological domains are interwoven across the lifespan. We demonstrate that individual behavioral domains are effectively associated with changes to multiple brain network systems. So whereas focussing on individual variables is helpful sometimes, it is not simply the case that individual behavioral variables affect individual brain areas or network systems. Opposite to that, the picture painted by our results show that many cognitive and emotional domains simultaneously vary across the lifespan with the changes in multiple brain networks. In sum, our results show major changes to behaviors and network structure that cannot be reduced to changes to any individual cognitive constructs. Instead, multiple cognitive and behavioral domains covary in ensemble with global changes to network structure. Future studies will be necessary to better describe the way the brain and behavior relationships reported here interact and how that drives aging in health and disease.

We demonstrate that brain and behavior show multiple global changes across the human lifespans that go well beyond one-to-one correlations between a single brain system or behavioral task. Several previous studies inspired this work, providing ideas for the novel contributions produced in this study. Early work [83] integrated functional brain networks and phenotypic data from a large dataset of healthy young adults [110]. A strong association between brain function and behavior was reported using CCA. Yet, age was not a significant result in the analysis as it was in ours, most likely due to the narrow age range in the Human Connectome Project sample used in that work compared to the CAN sample. Another relevant study used the CAN dataset, CCA, and generative models of brain functional activity and was first in reporting the effect of age on the association between cognitive performance and brain function [38]). The study focused on three functional networks [112,113] and 6 composite measures of behavior and showed that neuronal, not vasculature-related effects in resting state networks are associated with age. A third relevant study was first in using brain structure (a multimodal combination of diffusion-measures and cortical partial volume estimates) to show an association with various demographics, including age [84]. The current results go beyond previous work by (1) thoroughly cross-validating the CCA, (2) testing how well a hold-out variable is predicted by the primary canonical axis, and (3) using an RSA on the variable loadings from the cross-validated model to recover relationships between the variables themselves. These improvements provide a way for future work to glean more insight from the increasingly large and multimodal samples of variables utilized in modern neuroscience. Furthermore, we created open services shared on brainlife.io to allow other researchers to freely process new data utilizing our pipeline.

As neuroscience shifts towards data-driven approaches, data-fusion methods [114–116] such as CCA are likely to increase in popularity. Yet, the CCA approach used here is limited to two domains at a time. Other methods allow fusion across a higher number of data domains. For example, CCA has been generalized to higher-order models beyond 2 domains, similar to variations of Independent Component Analysis (joint- or linked-ICA; [117–119]. Imaging genetics is one of the fastest growing fields that are capitalizing the most on methods such as CCA especially requiring to combine multiple data domains [6,16,106,111,120,121]. These studies typically look at thousands of genetic markers from a single blood assay, similar to how a single MRI imaging session can be used to generate thousands of measures of brain structure and function. The benefit of methods such as CCA is due to the ability to model datasets with large numbers of variables from different domains (e.g. blood assays, behavior, genetics and neuroimaging). Yet, in addition to variations of CCA, ICA analyses have also been proposed to map across multiple variable domains, though Partial Least Squares (PLS; [82,122,123]) is perhaps the most common approach that allows mapping the combination of multiple data modalities into a single space [124,125]. Furthermore, the current work is also limited to linear interactions between data domains. Future explorations of the multivariate interactions might contribute additional novel insights [126,127]. Nonlinear interactions estimated through kernel embedding [128] or structural equation modelling [129] are particularly promising, but have not yet been applied to datasets similar to the one used here. Critically, one recent report has sparked interest by suggesting that adding nonlinear interactions to data modelling may provide negligible improvement over linear models when comparing brain-behavior interactions [130]. In selecting the best models (linear or nonlinear) might also depend on data preprocessing [131,132]. The effectiveness of data preprocessing may vary depending on the characteristics of the data domains. Future work will be required to determine what data domains will require linear or non-linear modelling approaches.

As neuroscience moves to population-level neuroscience, reproducible approaches to data management and analysis need to be embraced. Sharing data products is essential to implementing transparent, replicable, and reproducible brain research (23). Critical to the success of the next generation large-scale neuroscience methods will be to embrace new technology to ensure results reproducibility. This will require embracing methods for open science, data, and computational standards as well as modern computing infrastructure to lower the barriers of entry to proficient large-scale data methods (24–26). Our study follows up on the most recent trends in terms of large-scale datasets and computational approaches. But also the study is the first to embrace the most recent technology to support scientific reproducibility to clarify human aging. We used the recently developed, community-developed, and publicly-funded cloud computing platforms, brainlife.io. brainlife.io allows processing large amounts of data by tracking the provenance of each dataset and by linking the data-object on the platform with the reproducible web-services used to generate the data. brainlife.io addresses precisely the needs for replicability and reproducibility highlighted by the recent report by the U.S. National Academies (23). Indeed, to implement our study we developed more than ten new data analyses applications that are now available on brainlife.io for other investigators to reuse for new research or to replicate our results.

## Methods

### Data source

The behavior and neuroimaging data were accessed from the publicly distributed Cambridge center for Ageing and Neuroscience (CAN) dataset [90]. This large dataset provides cross-sectional data on 652 individuals. Of this total 623 had imaging data of interest to the current study (diffusion-weighted MRI and T1-weighted MRI). Out of a total of 623 individuals, 594 neuroimaging datasets were successfully processed and included in this present study. Quality control issues on the processed T1w data (FreeSurfer did not successfully segment the brain) motivated the exclusion of 26 subjects. **Supplemental Figure 2a** shows the dimensionality of the data (*n*=594).

### Behavioral data

Participants responded to a series of screening and demographic questionnaires and performed behavioral tasks and clinical tests [90]. See [90] for details on the tasks, tests and questionnaires. A total of 1,708 measurements per individual were acquired by the CAN consortium. Only behavior scores normalized by the CAN consortium were utilized for the current study, as a result, a total of 388 CAN-normalized scores were utilized for further analysis. All these measurements were initially subdivided into five of the primary behavioral domains: attention, memory, language, emotion, and motor [90]. For the current study, in addition to the five original behavioral domains, test scores collected by the CAN consortium measuring social engagement and clinical scores were also utilized and collected into a Social and Clinical behavioral domain for a total of 7 domains (**Figure 1a**). The social engagement measures consisted of 10 questions related to the frequency and mode of socialization of the individuals. Six clinical questionnaires were collected into the clinical domain: the MMSE, ACE-R, Wechsler Memory test, “Spot the Word”, the Cambridge 10MQ, and the PSQI Sleep Index [90,93].

### Neuroimaging Data Preprocessing

#### Anatomical MRI data preprocessing

A T1w and T2w anatomical scans were acquired for each individual. Both scans have a 1mm isotropic resolution (T1w: 3D MPRAGE, TR=2250ms, TE=2.99ms, TI=900ms; FA=9 deg; FOV=256×240×192mm; GRAPPA=2; TA=4mins 32s. T2w: 3D SPACE, TR=2800ms, TE=408ms, TI=900ms; FOV=256×256×192mm; GRAPPA=2; TA=4mins 30s). Both images were reoriented and aligned to the AC-PC plane with a 6 degree of freedom rigid alignment to the MNI template based on the Human Connectome Project (HCP) procedure (T1 reorientation; <u>bl.app.15</u>, T2 reorientation; <u>brainlife.app.114</u>, T1 alignment; <u>bl.app.99</u>, T2 alignment; <u>brainlife.app.116</u>). These AC-PC aligned images were passed to Freesurfer 6.0 [133], which reconstructs the white matter and pial surface based on the T1 image, using the T2 image for additional information to better estimate the surfaces (FreeSurfer; <u>bl.app.0</u>). The FreeSurfer output was used by the multi-atlas transfer tool (maTT; <u>bl.app.23</u>) to align individual subjects T1w anatomy files to the multimodal cortical brain atlas (MMP v1.0 Atlas [94]. The labels from the MMP atlas were used to build networks, see below.

#### Diffusion-weighted MRI data preprocessing

The diffusion-weighted MRI data (dMRI) contained a total of 60 directions collected across 2 b-value gradients (2D twice-refocused SE EPI, TR=9100ms, TE=104ms, TI=900ms; FOV=192×192mm; 66 axial slices, 2 mm isotropic; B0=0,1000/2000s/mm^2^, 30 unique directions per shell; TA=10mins 2s. Readout time 0.0684; echo spacing=0.72ms, EPI factor=96). Data were processed using MRtrix3 (<u>bl.app.68</u>; [96,134]. The following are the steps of the preprocessing procedure. The diffusion gradient alignment to the data was checked using a simple tracking procedure to find the orientation that produces the longest streamlines. After that, a PCA denoising procedure was performed to remove scanner noise not associated with the diffusion signal. Gibbs ringing, Eddy currents, inhomogeneity, and motion artifacts correction was performed. After all previous steps, an additional Rician-denoising was performed. The mean B0 image across (2 repeats) was extracted and utilized to register the dMRI to the AC-PC aligned T1-weighted image using Boundary Based Registration (BBR, [135]). The diffusion-weighting gradients were rotated to account for motion correction and AC-PC alignment. Finally, the diffusion-weighted image was resampled to a 1 mm isotropic.

#### Tractography generation

We developed an automated cloud computing service that we call Reproducible Anatomically Constrained Ensemble Tracking (RACE-Track; <u>bl.app.101</u>). RACE-Track combines Anatomically constrained tracking [98] with ensemble methods [97] and is containerized for public, reproducible access on the open cloud computing platform brainlife.io [89]. The preprocessed dMRI data was used to generate whole-brain tractography using RACE-Trac. Specifically, a constrained spherical deconvolution (CSD) response function was estimated for white matter, gray matter, and cerebrospinal fluid (CSF) as previously described [134,136]. Each respective response function was then used to estimate Fiber Orientation Distribution (FOD) maps for each tissue type. Following the ET approach, we fit the CSD model using multiple *L*_max_ (2, 4, and 6), this procedure created three FOD maps one per *L*_max_. The FOD maps were then passed to the iFOD2 tracking algorithm as previously described [98]. The T1w image was used to separate the brain tissue into its five types: 1) corticospinal fluid, 2) white matter, 3) gray matter, 4) subcortical gray matter and 5) pathological tissue. The image probability maps generated for each tissue were used for initiating and stopping tractography. More specifically, the gray-matter-white-matter interface was used as a seed mask for the streamlines, this approach has been demonstrated to provide a more complete coverage of tractography terminations on the cortex and subcortical gray matter structures [98]. Furthermore, the whole tractography procedure was repeated for both deterministic and probabilistic tracking, across the three *L*_max_ and five different maximum angles of curvature (5, 10, 20, 40, and 80º). A total of 600,000 streamlines was generated in each individual using this tracking procedure.

#### Construction of structural brain networks

Whole brain structural brain networks were built utilizing the multimodal cortical brain atlas (MMP v1.0; [94]). The MMP atlas was aligned to individual subjects’ brains with maTT (<u>bl.app.23</u>). The parcels of the MMP were used as nodes for the brain networks. The edges of the networks were defined as the density of connections between two parcels. Connection density was computed as the number of streamlines terminating in the two parcels divided by total number of voxels in the parcels [100]:

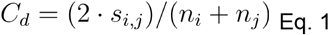

Where *C*_*d*_ is the estimated streamline density for a connection (network edge), *S*_*i,j*_ is the number of streamlines terminating in both regions, *n_i_* and *n_j_* are the number of voxels in each of the two brain regions. Networks matrices were created using Eq. 1 using a reproducible algorithm implemented on brainlife.io (<u>bl.app.121</u>). After that, the network matrices were thresholded by removing connections not present in at least half of the participants and then computing node degree for each individual network matrix using the Brain Connectivity Toolbox [91].

### Canonical correlation analysis and data preprocessing

#### Behavior features

The total of 388 behavioral variables (*m*_1_; **Supplemental Figure 2a**) were extracted from the CAN project. Data were prepared for modelling by applying a z-score transformation to each measurement across subjects [83]. Furthermore, variables with extreme outliers (more than three standard deviations from the population mean) or absent in at least half of subjects were eliminated from further analysis. Fifty-four of the behavioral variables were eliminated via normalization, bringing the total number of behavioral data utilized 388 to 334. These features were organized into a matrix (*D*_1_) composed of all behavioral (594 subjects, *n*, and 334 behavioral variables, *m*_1_). See **Supplemental Figure 2a** and **Equation 1**.

#### Network neuroscience features

Node degree was estimated using the streamline density networks generated described above for each subject [91]. This estimation procedure resulted in 376 node-degree measures per subject (*m*_2_). The upper diagonal of each network was unwinded to create a vector of features 1×70,500 in size [83]. These features were organized into a matrix (*D*_2_) composed of all brain data (594 subjects, *n*, and 70,500 node-degree estimates, *m*_2_). See **Supplemental Figure 2a** and **Equation 1**.

#### Data normalization

Both the brain and behavior datasets were normalized through the same process. First, the datasets were normalized by removing the mean of each feature across subjects and dividing by the standard deviation across subjects. Next, nuisance variables were regressed from each dataset to remove their influence on the data. These variables include a subjects’ height, weight, gender, heart rate and systolic and diastolic blood pressure. Behavior data was also rotated to the nearest positive definite pairwise covariance matrix to better account for correlation between variables [83]. This step was not required and not performed on the brain network data. Principal component analysis (PCA) was performed on *D*_1_ and *D*_2_ separately to reduce the number of dimensions in the data. PCA generated two eigenvector matrices, *E*_1_ and *E*_2_, for *D*_1_ and *D*_2_, respectively (see **Supplemental Figure 2b** and **Equation 1**). Because the choice of number of principal components (PCs) is arbitrary, we explored a wide range of PCs (2-100) and selected the PC number for *D*_1_ and *D*_2_ that best predicted the individual subjects’ age (see below for more details).

#### Canonical correlation analysis (CCA)

We used *E*_1_ and *E*_2_ to perform canonical correlation analysis (see **Supplemental Figure 2c**). The CCA algorithm estimates the linear combination of variables from each dataset that maximizes the correlation between the datasets

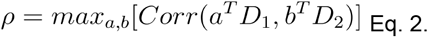

where ρ is the correlation between the datasets being maximized, *D*_1_ and *D*_2_ are the normalized datasets, and *a* and *b* are the weighting matrices that linearly combine the data variables into the canonical scores. The CCA analysis returns two matrices, *F*_1_ and *F*_2_, containing the canonical factor scores for each subject to each of the estimated canonical factors. The factor scores describe how much each subject contributes to each of the latent factors. These factor scores are used plotting the canonical correlation axes, or canonical axes (CA, **Figure 2**, **Supplemental Figure 2**, red). We use the combination of the first factor scores from each dataset to estimate the correlation with age.

To better interpret the contribution of individual variables within the CCA factors, variable loadings are reconstructed from the normalized data and the canonical factors. The loading values, *L*_1_ and *L*_2_ for brain and behavior respectively, describe the strength of the variables contribution to the axis. The loading matrices’ dimensions are the number of variables in the dataset by the number of estimated canonical factors. Each variable has a loading for every factor. The loadings are recovered for a variable by correlating the original normalized variables for every subject with the corresponding factor scores. This process is repeated and every variable is correlated with a factor to create the variable loadings for a canonical axis. By correlating the normalized variable observations (each column of matrix *D*) to each factor score (each column of matrix *F*) (**Supplemental Figure 2**) for every canonical factor estimated in both datasets, we generate canonical loadings (*L*) for every variable and every factor. For example, a specific behavioral score could be highly correlated with a canonical factor, indicating that the behavior is highly predictive of changes in this factor. When this is done for every axis, a complete matrix of variable loadings is recovered. The highest variable loadings for a canonical axis indicate the variables that contribute most to the positive end of the canonical factor, while the lowest variable loadings indicate the variables that contribute most to the negative end of the canonical factor. This is reported in Figure 3. The variable loadings are the typical way of inspecting how variables within a CCA analysis interact.

#### Correlation with Age

To determine the association of a canonical axis with age, we used a multi-way correlation with the first canonical factors to determine an r coefficient between the canonical axis and age. Age was predicted by

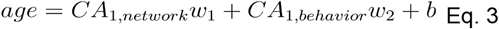

where *age* is the age of each subject, *CA_1, network_* is every subjects score in the first column of the brain factors (*F*_1_), *CA_1,behavior_* is every subjects score in the first column of the behavior factors (*F*_2_), *w*_1_ and *w*_2_ are the beta estimates of the model, and is the intercept. This is an ordinary least squares linear regression using the canonical scores of the behavior and brain from the first factor predicting age. To determine the correlation between the factors and age, we take the square root of the R2 from the fit model to estimate the correlation coefficient between the combination of factors and age. This is equivalent to the correlation with age along the canonical axis of the estimated factors. To determine the significance of the correlation of age to the canonical axis, a bootstrap test of 10,000 permutations was performed by randomly sorting the rows with replacement of *CA_1, network_* and *CA_1,behavior_* to determine a null prediction. This allowed us to determine if the observed correlation with age was significantly different from zero. To estimate the standard error of the estimate, a separate 10,000 permutation bootstrap was estimated, sampling with replacement both factors and age. The resampled estimates of age were used to create a rempled mean and standard error of the correlation between the canonical factors and age.

#### Parameter tuning

The goal of this analysis is to maximize the correlation with age. However, the principal components analysis performed as part of the preparation of the data for the CCA will combine variables differently depending on the number of components requested (**Supplemental Figure 2f**, **g**, **h**). Further, the number of PCA components indicates the number of canonical factors that can be estimated. Typically, a number such as 20 or 100 is selected by the researcher. However, there is reason to believe that the number of requested components will impact the final correlation with age (**Supplemental Figure 2f**, **g**, **h**). To test this we tuned the number of PCA components using a grid search, selecting the number of principal components based on the strength of the resulting correlation with age. This hyperparameter sweep found the correlation with age on the primary canonical factors for every combination of PCA components between 2 and 100 on the input datasets. See Supplemental Figure **2e, d** for the results from the search space. We selected 38 principal components for the brain axis and 40 principal components for the behavior axis because this combination of principal components resulted in CCA factors with the highest correlation with age. Nearby parameter spaces had similar levels of correlation with age. Importantly, the most common combinations of parameters, [20×20] and [100×100], had a significantly worse correlation with age than the model with the optimal parameters (**Supplemental Figure 2f**, **g**, **h**). This is further supported by inspecting the variable loadings of the models that performed worse than the tuned solution. The variable loadings in the suboptimal models had different combinations of variables as the strongest and weakest loadings. This difference indicates that the CCA used a different combination of variables to form the canonical factors in these models. These different factors are less effective at predicting age. This indicates that the selection of the number of principal components for this analysis is important in tuning an accurate model.

#### Model selection

All factors and loadings were cross-validated using a repeated 5-fold process to better ensure the generalizability of this model to the larger population (**Supplemental Figure 2e**). A repeated k-fold process is conceptually similar to a typical k-fold approach. A standard k-fold approach models data by training the parameters on a majority portion of the subjects and then applying the fit parameters to a holdout set of subjects to observe the generalizability of the trained parameters to new samples from the same population. The two key differences between the repeated k-fold process and a standard k-fold process is that 1) in a repeated k-fold process all observations end up with a held out estimate and 2) the training/test process is repeated many times to estimate a distribution of hold out parameters. The central tendency of the hold out parameter’s distribution is taken for each of the subject factors and variable loadings. For example, a standard k-fold cross-validation would divide the subjects evenly into an 80-20 train/test split. The CCA would be fit, or trained, on 80% of the subjects. The trained parameters from 80% of the subjects would be applied to the 20% holdout. The canonical factors and variable loadings are compared between the train and test data to determine how well the model generalizes to the population.

To perform the repeated k-fold process, the dataset was first split into 5 equal groups of subjects. Then the CCA was fit 5 times using each combination of 4 of the groups to train the model, predicting the holdout estimates for each of the splits. This creates a full set of hold out estimates for every subject’s factors (*F*) and loadings (*L*). This process is repeated 15,000 times, with each iteration creating a new random split of subjects into 5 groups before repeating the train/test procedure. This creates a distribution of trained factors and loadings for every parameter in the model. The median value of these distributions indicates the most probable trained value for the parameter. Additionally, the central tendency of these distributions is observed to quantify the standard error of the estimates. These median values of the permuted hold-out distributions are the factors and loadings used in evaluating the CCA (the factors values in Fig 1, the loading values in Fig 2). The repeated k-fold cross-validation approach allows the full dataset to be evaluated in the results, as each subject’s factor scores and each variable’s loadings are estimated from a model using unique observations to predict their estimate. This approach was necessary because the modeling of data with a CCA does not generalize to new observations well. The estimation of the CCA is based largely on deterministic matrix algebra, making the final results very sensitive to small perturbations in the input data. Selecting new training-test splits from the observations in a single k-fold cross-validation approach often resulted in different conclusions, while this repeated k-fold cross-validation consistently produces the same model estimates.

Additionally, this cross-validation approach is robust to our data being evenly sampled across our outcome of interest, age. Any one random sample of subjects from this dataset would be likely to bias the training or prediction of a particular age. Further, it is not possible to use age as part of the identification of training subjects while still being able to use it as an unbiased outcome. By cross-validating across many random k-fold splits we are able to cross-validate our model without risking an unintended bias with age that may be present in any one split.

#### Bootstrap Tests

A bootstrap method was used to test the statistical significance of the correlations and loadings within the CCA models (*M*_0_, *M*_1_ and *M*_2_). The bootstrap tests were implemented by assuming a null hypothesis of “no correlation between the two variables (or parameters sets).” For each of the various tests described in the results section, a sampling distribution of the mean was generated by resampling the data (or model parameters) with replacement. For each bootstrap test, 10,000 samples of the data (or model parameters) were generated. The probability of the empirically observed correlation given the sampling distribution was reported (*p*). Gray shading in **Supplemental Figure 2i**, **j** show examples of the resampling distributions generated by bootstrap under the null hypothesis of no correlation between variables.

#### Building the average contribution of brain networks and behavior domains as plots and word clouds

To summarize the brain networks and behavior domains that most strongly contribute to *M_1_*, the variables within each dataset were averaged within predetermined modules and summarized with a scatter plot and word cloud plot. The module for the brain network loadings was built by assigning the labels from the MMP (v1.0) atlas to a set of established functional network labels from [93], referred to as Y_2011_. Each of the 376 nodes in our networks was assigned to one of the seven functional networks in Y_2011_; Visual, Somatomotor, Limbic, Ventral attention, Dorsal Attention, Frontoparietal, Default Mode (DMN). In addition, the hippocampus, amygdala were kept separated and all the remaining subcortical structures were combined and reported as subcortical (i.e., pallidum, putamen, accumbens, thalamus, and caudate). This process generated a total of ten labels (words) representing the established functional brain networks of Y_2011_, referred to as modules. The modules for the behavioral domain were determined from the CAN domains that the tasks were chosen to sample across (Memory, Language, Emotion, Attention, Motor, Clinical Scores, and Social). First, the individual variable loadings were taken from the cross-validated model. Next, the variables from each axis were combined within each module by averaging together all the positive values into a positive average and the negative values into a negative average. By creating separate averages for the positive and negative loadings, the prevalence of the modules (brain network or behavior) on each end of the axis was assessed. The error bars were computed by taking the average of the respective variables’ standard errors from the cross-validated model. The left-hand size plots in **Figure 3c** and **Figure 3d** display the average variable loading within each network or behavioral module as a pair of points. The positive average (*blue*) and negative average (*red*) contribution of variables within the respective module are displayed as a pair of columns. Error bars show 2 units of the average standard error estimated by cross-validation. The right hand side plots in **Figure 3c** and **Figure 3d** show word-cloud representations of the variable modules. These plots were created using the same average values (**Figure 3c** and **d** *left hand side*) scaling the font size by sum of the modules loadings. More specifically, the font size of each word was scaled by the average of the loadings within each module. The larger the font of the word, the larger the average for positive values (*blue*) or lower for negative values (*red*). For example, on the behavioral axis **(Figure 2a**, *y-axis*), on average all the memory tasks contributed a lower score than the emotion tasks, hence the font size for memory is smaller than that for emotion in **Figure 3d**.

#### Computing the rich club of the network and its comparison to the CCA

In order to compare the CCA model to the rich club (RC; [103]), a global network was constructed by averaging individual networks across all subjects. A standard consensus threshold was applied after averaging by zeroing any connection not present in a simple majority of individuals (i.e., a connection was kept in the average matrix if the connection was available in more than 50% of the total number of subjects). Next, node degree was estimated on this averaged network using the BCT [91]. The node degree estimates were normalized by subtracting the minimum and dividing by the maximum. A threshold was then applied to the node degree to identify the RC. We used results from previous work [103] and set the threshold of 14% to separate RC from periphery. This means that the top 14% of the nodes with the highest normalized node degree were assigned to RC. More specifically, 54 of 376 regions in the network were assigned to the rich club the remaining 322 to the periphery –see color-coded ball-stick representations in **Figure 4**. The size of the nodes in **Figure 4** were scaled by the magnitude of the CCA loading of each node.

After identifying the RC and periphery, the association between RC/Periphery and the CCA was investigated. The spearman rank correlation was estimated between the node degree in each node and CCA loadings. A bootstrap test was used to estimate the mean, standard error, and p-values reported in **Results**. To visualize the correlations, a scatter plot comparing the CCA variable loadings and the participation coefficient of the RC is shown in **Supplemental Figure 4a**. The points are color coded for the rich club (*blue*) and periphery (*gray*). The ordinate of **Supplemental Figure 4a** is the RC participation coefficient for each node. The participation coefficient was computed as the ratio of the number of connections from that node to other nodes in the same group (RC or periphery) to the number of connections from that node to the other groups [103]. The full list of CCA loadings in relation to the participation coefficient was displayed (**Supplemental Figure 4b**) to show that the majority of RC connections also have relatively high loadings within the CCA as well.

### Representational Similarity Analysis (RSA)

We used Representational Similarity Analysis (RSA, [92,139]) to create a fingerprint of how the loadings of the variables in one data domain (brain) are associated with the loadings of the variables in the other data domain (behavior). We define factors as the estimated latent factors from the CCA analysis. The number of factors estimated is determined by the smallest number of tuned PCA parameters used during parameter tuning. Each variable has a loading for the factors estimated. We use a dissimilarity measure estimated between pairs of variable loadings to inspect how the variables interact with each other across all estimated factors. This allows us to observe how the individual observations within the domains of our CCA model, brain and behavior, interact with each other as they form the primary axes for all factors in the analysis. Typically, the extent of our inspection of the CCA model is using the cross-validated variable loadings ranked by their magnitude to indicate the strongest contributing variables from a particular domain (**Figure 2**). However, there is currently no way to recover how the individual variables interact with each other. While the CCA determines an overall interaction between the separate datasets and the variables that most contribute to each factor, there was previously no description of how the individual elements interact with one another in this model. Access to this information would prove particularly useful to determine what relationships within the data may be capitalized on to estimate the canonical factors, as there are many established brain-behavior interactions within this dataset.

The dissimilarity matrix (*S*) is positive and block symmetric, with three distinct components representing the network-to-network (*ntn*), behavior-to-behavior (*btb*) and brain-to-behavior (*btn*) dissimilarity. After building *S*, the elements of the matrix were sorted and organized into meaningful brain and behavioral domains. This organization allowed us to interpret values of *S* in terms of functional domains for both brain and behavior instead of individual regions or tasks. Multiple individual brain regions in *θ*_B_ were sorted and uniquely assigned into the Y_2011_ labels ([93]; https://github.com/bcmcpher/cca_aging/blob/master/HCP_to_Yeo_assignment.tsv). The behavioral measurements in *θ*_b_ were mapped onto the seven major behavioral domains and averaged (Social, Emotional, Attention, Memory, Clinical, Language, and Motor). Results show that the CCA captures patterns of covariation between the two domains (brain and behavior). For example, the emotion domain is highly similar (low dissimilarity) both in the brain and behavior between the brain and behavioral contributions. Also, attention and memory are highly similar (low dissimilarity) as well. To contrast, these results to the normalized data, see **Supplemental Figure 4i-k** that shows the difference in range between the dissimilarity of the observed variables (x-axis) to the CCA estimates (y-axis).

To build the dissimilarity matrix between all variables used in the analysis, we first use the variable loadings reconstructed from the cross-validated factors. To construct the RSA, we take the dissimilarity measure between every pair of variables using their recovered loadings for all factors. This is taken across both *L*_1_ and *L*_2_ (the loading matrices for brain and behavior, respectively). Each row represents the canonical loadings for every canonical factor for the variable and each column is the order of the estimated canonical factors. By taking the dissimilarity measure between each row, we get a measure of how similarly any two variables behave across all the estimated factors. The dissimilarity is able to be taken across domains, meaning how the brain and behavior variables interact within their respective factors can be inspected directly.

The first factor of the CCA represents the factor structure of each domain that creates the strongest correlation between the datasets, while each subsequent factor represents the next orthogonal configuration that has the next strongest correlation. By comparing the variable loadings of each factor between two variables we can represent how the variables interact with each other across every estimated canonical factor. We use this full vector of estimated factor scores to determine the distance by calculating the dissimilarity between every unique pair of variables. A dissimilarity measure:

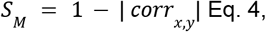

where x and y are any pair of variable loadings in an [n x 1] vector. Each vector consists of the *n* loadings for each canonical factor estimated in the model for each variable. The correlation between these vectors would represent how similarly the variables contribute to each of the estimated canonical factors. Every unique combination of x and y created a symmetric matrix of size [710 × 710] with the respective dissimilarity value between every variable estimated. The values represent how dissimilar any pair of variables behave across every estimated factor. The higher the score between the variables, the more independently they interact across all the estimated canonical factors. Likewise, the lower the score between the variables, the more similarly they behave across factors. To summarize this information, we sorted these measures into established behavior domains and brain networks. This better reveals how variables within a specific domain behave with variables from another domain. From this sorted matrix, we estimate the modules by averaging each value within a module into a single value. This creates a matrix of size [modules x modules] with the mean and standard deviation of dissimilarity values for every variable within that group. These values represent on average how the behavior domains and brain networks interact with each other across all the factors. However, the most important interactions for our purpose is the interaction between the brain-behavior averages, or how well the behavior domains correspond to the structural connectivity within established functional networks (**Figure 4e**).

To do so, we computed the loadings by correlating the original normalized data and the CCA factors for each canonical correlate estimated (columns of *D_1,2_* with columns of *F_1,2_*, respectively). Two matrices were generated with this approach, one for the behavioral and one for the brain network loadings (*L_1_* and *L_2_*). These loadings were then stacked into a matrix where each row is a variable and each column is the corresponding canonical axis (**Supplemental Figure 4a**) and used to compute the dissimilarity matrix between all brain regions and behavioral tasks. This creates a symmetric matrix where the pairwise dissimilarity distance between all rows of the stacked matrices was computed using Eq. 4 (**Figure 3a**; [92]). The axes are the size of the number of variables included in the model and a value for each pair of variables indicating how similarly they behave across all canonical factors.

#### Chord Plot

To better visualize the association between brain and behavior we transformed the brain-behavior dissimilarity value back to correlations: *r* = 1 − *S*_*M*_. These values were then used for a modified chord plot visualization. For this plot, the *r* values were normalized by rescaling them to better show the brain-behavior interactions. First, *r* was normalized between 0 and 1. Second, *r* values below the top 75th percentile were eliminated (**Supplementary Figure 4h-j**). Third, *r* was multiplied by 1,000 and squared to visually emphasize any difference in values. The chord plots represent the strength of the association between the CCA loadings of the brain networks and behavioral domains. Two aspects of this plot informa our analysis. (A) Each wedge of the chord shows the total association in a domain; the larger the wedge for a particular domain the larger the contribution to the CCA loadings. (B) The thicker the bands connecting a behavioral domain and a brain network, the stronger the association between the loadings in those variables.

#### Evidencing the effect of age to the CCA by computing the difference in RSA between *M*_1_ **and** *M*_2_

To determine the impact of age on the brain-behavior relationship, we repeated the whole cross-validated CCA procedure, but first age was regressed along with the other covariates before the CCA was performed, referred to as *M_2_* in the text. This effectively removes the association with age from the subsequent analyses as described in Results. When the RSA analysis is repeated with *M_2_*, the resulting dissimilarity modules reflect the brain-behavior interaction with the variability of age removed. What is left represents the difference between models with and without age. By taking the difference between the *S_1_* and *S_2_* we are able to show the impact of age on the brain-behavior relationship (*S_d_*). The difference between these modules is the impact of age on the connection between these scores.

## Acknowledgments

This research was supported by NSF IIS-1912270, NSF IIS-1636893, NSF BCS-1734853, Microsoft Faculty Fellowship to F.P., NIH 5T32MH103213-05 to William Hetrick. We thank Soichi Hayashi, Brad Caron and Josh Faskowitz for contributing to the development of brainlife.io, Dan Stanzione, William. J. Allen, for support at TACC supercomputing center and Craig Stewart, Robert Henschel, David Hancock and Jeremy Fischer for support with jetstream-cloud.org (NSF ACI-1445604).

## Notes

### Competing Interest Statement

The authors have declared no competing interest.

### Summary of Updates

Revised draft based on reviewer feedback for R&R.

## References

1. Nations U, United Nations. World Population Ageing 2019 Highlights. 2019. doi:10.18356/9df3caed-en

2. Rich PB, Barry N. Health-Care Economics and the Impact of Aging on Rising Health-Care Costs. Geriatric Trauma and Critical Care. 2017. pp. 99–105. doi:10.1007/978-3-319-48687-1_11

3. Lustig C, Shah P, Seidler R, Reuter-Lorenz PA. Aging, Training, and the Brain: A Review and Future Directions. Neuropsychology Review. 2009. pp. 504–522. doi:10.1007/s11065-009-9119-9

4. Deary IJ, Corley J, Gow AJ, Harris SE, Houlihan LM, Marioni RE, et al. Age-associated cognitive decline. Br Med Bull. 2009;92: 135–152.

5. Niccoli T, Partridge L. Ageing as a Risk Factor for Disease. Current Biology. 2012. pp. R741–R752. doi:10.1016/j.cub.2012.07.024

6. Risacher SL, Saykin AJ. Neuroimaging in aging and neurologic diseases. Handbook of Clinical Neurology. 2019. pp. 191–227. doi:10.1016/b978-0-12-804766-8.00012-1

7. Harada CN, Natelson Love MC, Triebel KL. Normal Cognitive Aging. Clinics in Geriatric Medicine. 2013. pp. 737–752. doi:10.1016/j.cger.2013.07.002

8. Wykes T, Haro JM, Belli SR, Obradors-Tarragó C, Arango C, Ayuso-Mateos JL, et al. Mental health research priorities for Europe. Lancet Psychiatry. 2015;2: 1036–1042.

9. DiLuca M, Olesen J. The cost of brain diseases: a burden or a challenge? Neuron. 2014;82: 1205–1208.

10. Gustavsson A, Svensson M, Jacobi F, Allgulander C, Alonso J, Beghi E, et al. Cost of disorders of the brain in Europe 2010. Eur Neuropsychopharmacol. 2011;21: 718–779.

11. Andlin-Sobocki P, Jönsson B, Wittchen H-U, Olesen J. Cost of disorders of the brain in Europe. Eur J Neurol. 2005;12: 1–27.

12. Wittchen HU, Jacobi F, Rehm J, Gustavsson A, Svensson M, Jönsson B, et al. The size and burden of mental disorders and other disorders of the brain in Europe 2010. Eur Neuropsychopharmacol. 2011;21: 655–679.

13. Taylor WD, Raji C, Wang L, Lavretsky H. Can We Change the Inevitable? Ameliorating Brain Aging and Cognitive Decline. The American Journal of Geriatric Psychiatry. 2016. p. S7. doi:10.1016/j.jagp.2016.01.008

14. Vinke EJ, de Groot M, Venkatraghavan V, Klein S, Niessen WJ, Ikram MA, et al. Trajectories of imaging markers in brain aging: the Rotterdam Study. Neurobiol Aging. 2018;71: 32–40.

15. Beaton D, Rieck JR, Alhazmi F, Abdi H, ADNI. Multivariate genotypic analyses that identify specific genotypes to characterize disease and control groups in ADNI. doi:10.1101/235945

16. Veitch DP, Weiner MW, Aisen PS, Beckett LA, Cairns NJ, Green RC, et al. Understanding disease progression and improving Alzheimer’s disease clinical trials: Recent highlights from the Alzheimer’s Disease Neuroimaging Initiative. Alzheimer’s & Dementia. 2019. pp. 106–152. doi:10.1016/j.jalz.2018.08.005

17. Svaldi DO, Goñi J, Amico E, Abbas K, Charanya M, West JD, et al. IC-P-032: IMPROVING PREDICTION OF COGNITIVE OUTCOMES FROM FUNCTIONAL CONNECTIVITY IN ALZHEIMER’S DISEASE. Alzheimer’s & Dementia. 2019. pp. P38–P39. doi:10.1016/j.jalz.2019.06.4194

18. Park DC, Reuter-Lorenz P. The adaptive brain: aging and neurocognitive scaffolding. Annu Rev Psychol. 2009;60: 173–196.

19. Cepeda NJ, Kramer AF, Gonzalez de Sather JC. Changes in executive control across the life span: examination of task-switching performance. Dev Psychol. 2001;37: 715–730.

20. Amunts J, Camilleri JA, Eickhoff SB, Heim S, Weis S. Executive functions predict verbal fluency scores in healthy participants. Sci Rep. 2020;10: 11141.

21. Salthouse TA. Theoretical Perspectives on Cognitive Aging. 2016. doi:10.4324/9781315785363

22. Salthouse TA. Trajectories of normal cognitive aging. Psychology and Aging. 2019. pp. 17–24. doi:10.1037/pag0000288

23. Cabeza R, Nyberg L, Park D. Cognitive Neuroscience of Aging: Linking Cognitive and Cerebral Aging. Oxford University Press; 2009.

24. Kramer AF, Hahn S, Gopher D. Task coordination and aging: explorations of executive control processes in the task switching paradigm. Acta Psychologica. 1999. pp. 339–378. doi:10.1016/s0001-6918(99)00011-6

25. Salthouse TA. What and When of Cognitive Aging. Current Directions in Psychological Science. 2004. pp. 140–144. doi:10.1111/j.0963-7214.2004.00293.x

26. Ratcliff R, Thapar A, Gomez P, McKoon G. A diffusion model analysis of the effects of aging in the lexical-decision task. Psychol Aging. 2004;19: 278–289.

27. Thapar A, Ratcliff R, McKoon G. A diffusion model analysis of the effects of aging on letter discrimination. Psychol Aging. 2003;18: 415–429.

28. Lisman J, Buzsáki G, Eichenbaum H, Nadel L, Ranganath C, David Redish A. Publisher Correction: Viewpoints: how the hippocampus contributes to memory, navigation and cognition. Nature Neuroscience. 2018. pp. 1018–1018. doi:10.1038/s41593-017-0034-8

29. Nyberg L, Pudas S. Successful Memory Aging. Annu Rev Psychol. 2019;70: 219–243.

30. Lu Z-L, Neuse J, Madigan S, Dosher BA. Fast decay of iconic memory in observers with mild cognitive impairments. Proc Natl Acad Sci U S A. 2005;102: 1797–1802.

31. Sekuler R, Hutman LP. Spatial Vision and Aging. I: Contrast Sensitivity. Journal of Gerontology. 1980. pp. 692–699. doi:10.1093/geronj/35.5.692

32. Bennett PJ, Sekuler R, Sekuler AB. The effects of aging on motion detection and direction identification. Vision Research. 2007. pp. 799–809. doi:10.1016/j.visres.2007.01.001

33. Ratcliff R, Thapar A, McKoon G. A diffusion model analysis of the effects of aging on brightness discrimination. Percept Psychophys. 2003;65: 523–535.

34. Samanez-Larkin GR, Robertson ER, Mikels JA, Carstensen LL, Gotlib IH. Selective attention to emotion in the aging brain. Psychol Aging. 2009;24: 519–529.

35. Carstensen LL, Turan B, Scheibe S, Ram N, Ersner-Hershfield H, Samanez-Larkin GR, et al. Emotional experience improves with age: evidence based on over 10 years of experience sampling. Psychol Aging. 2011;26: 21–33.

36. Mok RM, Myers NE, Wallis G, Nobre AC. Behavioral and Neural Markers of Flexible Attention over Working Memory in Aging. Cerebral Cortex. 2016. pp. 1831–1842. doi:10.1093/cercor/bhw011

37. Gaetz W, Roberts TPL, Singh KD, Muthukumaraswamy SD. Functional and structural correlates of the aging brain: Relating visual cortex (V1) gamma band responses to age‐related structural change. Human Brain Mapping. 2012. pp. 2035–2046. doi:10.1002/hbm.21339

38. Tsvetanov KA, Henson RNA, Tyler LK, Razi A, Geerligs L, Ham TE, et al. Extrinsic and Intrinsic Brain Network Connectivity Maintains Cognition across the Lifespan Despite Accelerated Decay of Regional Brain Activation. J Neurosci. 2016;36: 3115–3126.

39. Grady C. The cognitive neuroscience of ageing. Nature Reviews Neuroscience. 2012. pp. 491–505. doi:10.1038/nrn3256

40. Grady CL, Protzner AB, Kovacevic N, Strother SC, Afshin-Pour B, Wojtowicz M, et al. A multivariate analysis of age-related differences in default mode and task-positive networks across multiple cognitive domains. Cereb Cortex. 2010;20: 1432–1447.

41. Fjell AM, Walhovd KB, Fennema-Notestine C, McEvoy LK, Hagler DJ, Holland D, et al. One-year brain atrophy evident in healthy aging. J Neurosci. 2009;29: 15223–15231.

42. Johnson SC, Saykin AJ, Baxter LC, Flashman LA, Santulli RB, McAllister TW, et al. The relationship between fMRI activation and cerebral atrophy: comparison of normal aging and alzheimer disease. Neuroimage. 2000;11: 179–187.

43. Apostolova LG, Green AE, Babakchanian S, Hwang KS, Chou Y-Y, Toga AW, et al. Hippocampal atrophy and ventricular enlargement in normal aging, mild cognitive impairment (MCI), and Alzheimer Disease. Alzheimer Dis Assoc Disord. 2012;26: 17–27.

44. Driscoll I, Hamilton DA, Petropoulos H, Yeo RA, Brooks WM, Baumgartner RN, et al. The aging hippocampus: cognitive, biochemical and structural findings. Cereb Cortex. 2003;13: 1344–1351.

45. Habes M, Janowitz D, Erus G, Toledo JB, Resnick SM, Doshi J, et al. Advanced brain aging: relationship with epidemiologic and genetic risk factors, and overlap with Alzheimer disease atrophy patterns. Transl Psychiatry. 2016;6: e775.

46. Salat DH, Buckner RL, Snyder AZ, Greve DN, Desikan RSR, Busa E, et al. Thinning of the cerebral cortex in aging. Cereb Cortex. 2004;14: 721–730.

47. Maillet D, Rajah MN. Association between prefrontal activity and volume change in prefrontal and medial temporal lobes in aging and dementia: a review. Ageing Res Rev. 2013;12: 479–489.

48. Paxton JL, Barch DM, Racine CA, Braver TS. Cognitive control, goal maintenance, and prefrontal function in healthy aging. Cereb Cortex. 2008;18: 1010–1028.

49. Huettel SA, Singerman JD, McCarthy G. The effects of aging upon the hemodynamic response measured by functional MRI. Neuroimage. 2001;13: 161–175.

50. D’Esposito M, Zarahn E, Aguirre GK, Rypma B. The effect of normal aging on the coupling of neural activity to the bold hemodynamic response. Neuroimage. 1999;10: 6–14.

51. Buckner RL, Snyder AZ, Sanders AL, Raichle ME, Morris JC. Functional Brain Imaging of Young, Nondemented, and Demented Older Adults. Journal of Cognitive Neuroscience. 2000. pp. 24–34. doi:10.1162/089892900564046

52. Li H-J, Hou X-H, Liu H-H, Yue C-L, Lu G-M, Zuo X-N. Putting age-related task activation into large-scale brain networks: A meta-analysis of 114 fMRI studies on healthy aging. Neurosci Biobehav Rev. 2015;57: 156–174.

53. Reuter-Lorenz PA, Lustig C. Brain aging: reorganizing discoveries about the aging mind. Curr Opin Neurobiol. 2005;15: 245–251.

54. Milham MP, Erickson KI, Banich MT, Kramer AF, Webb A, Wszalek T, et al. Attentional Control in the Aging Brain: Insights from an fMRI Study of the Stroop Task. Brain and Cognition. 2002. pp. 277–296. doi:10.1006/brcg.2001.1501

55. Gordon BA, Zacks JM, Blazey T, Benzinger TLS, Morris JC, Fagan AM, et al. Task-evoked fMRI changes in attention networks are associated with preclinical Alzheimer’s disease biomarkers. Neurobiology of Aging. 2015. pp. 1771–1779. doi:10.1016/j.neurobiolaging.2015.01.019

56. Gazzaley A, Cooney JW, Rissman J, D’Esposito M. Top-down suppression deficit underlies working memory impairment in normal aging. Nat Neurosci. 2005;8: 1298–1300.

57. Yeatman JD, Wandell BA, Mezer AA. Lifespan maturation and degeneration of human brain white matter. Nature Communications. 2014. doi:10.1038/ncomms5932

58. Thomason ME, Thompson PM. Diffusion Imaging, White Matter, and Psychopathology. Annual Review of Clinical Psychology. 2011. pp. 63–85. doi:10.1146/annurev-clinpsy-032210-104507

59. Reisberg B, Franssen EH, Souren LEM, Auer SR, Akram I, Kenowsky S. Evidence and mechanisms of retrogenesis in Alzheimer’s and other dementias: Management and treatment import. American Journal of Alzheimer’s Disease & Other Dementiasr. 2002. pp. 202–212. doi:10.1177/153331750201700411

60. Cox SR, Ritchie SJ, Tucker-Drob EM, Liewald DC, Hagenaars SP, Davies G, et al. Ageing and brain white matter structure in 3,513 UK Biobank participants. Nature Communications. 2016. doi:10.1038/ncomms13629

61. Xu T, Yang Z, Jiang L, Xing X-X, Zuo X-N. A Connectome Computation System for discovery science of brain. Science Bulletin. 2015. pp. 86–95. doi:10.1007/s11434-014-0698-3

62. Zuo X-N, He Y, Betzel RF, Colcombe S, Sporns O, Milham MP. Human Connectomics across the Life Span. Trends Cogn Sci. 2017;21: 32–45.

63. Salthouse TA. Neuroanatomical substrates of age-related cognitive decline. Psychol Bull. 2011;137: 753–784.

64. Wandell BA. Clarifying Human White Matter. Annu Rev Neurosci. 2016;39: 103–128.

65. Gutchess A. Plasticity of the aging brain: new directions in cognitive neuroscience. Science. 2014;346: 579–582.

66. Oschwald J, Guye S, Liem F, Rast P, Willis S, Röcke C, et al. Brain structure and cognitive ability in healthy aging: a review on longitudinal correlated change. Rev Neurosci. 2019;31: 1–57.

67. Cole JH, Franke K. Predicting Age Using Neuroimaging: Innovative Brain Ageing Biomarkers. Trends Neurosci. 2017;40: 681–690.

68. Cole J, Franke K, Cherbuin N. Quantification of the biological age of the brain using neuroimaging. doi:10.31219/osf.io/3b6zu

69. Bishop NA, Lu T, Yankner BA. Neural mechanisms of ageing and cognitive decline. Nature. 2010. pp. 529–535. doi:10.1038/nature08983

70. Turner BM, Forstmann BU, Love BC, Palmeri TJ, Van Maanen L. Approaches to Analysis in Model-based Cognitive Neuroscience. J Math Psychol. 2017;76: 65–79.

71. Falk EB, Hyde LW, Mitchell C, Faul J, Gonzalez R, Heitzeg MM, et al. What is a representative brain? Neuroscience meets population science. Proc Natl Acad Sci U S A. 2013;110: 17615–17622.

72. Poldrack RA. Mapping Mental Function to Brain Structure: How Can Cognitive Neuroimaging Succeed? Perspect Psychol Sci. 2010;5: 753–761.

73. Bassett DS, Sporns O. Network neuroscience. Nat Neurosci. 2017;20: 353–364.

74. van den Heuvel MP, Sporns O. A cross-disorder connectome landscape of brain dysconnectivity. Nat Rev Neurosci. 2019;20: 435–446.

75. Petersen SE, Sporns O. Brain Networks and Cognitive Architectures. Neuron. 2015. pp. 207–219. doi:10.1016/j.neuron.2015.09.027

76. Braun U, Schäfer A, Walter H, Erk S, Romanczuk-Seiferth N, Haddad L, et al. Dynamic reconfiguration of frontal brain networks during executive cognition in humans. Proc Natl Acad Sci U S A. 2015;112: 11678–11683.

77. Khambhati AN, Mattar MG, Wymbs NF, Grafton ST, Bassett DS. Beyond modularity: Fine-scale mechanisms and rules for brain network reconfiguration. Neuroimage. 2018;166: 385–399.

78. Rutter LA, Vahia IV, Forester BP, Ressler KJ, Germine L. Heterogeneous Indicators of Cognitive Performance and Performance Variability Across the Lifespan. Front Aging Neurosci. 2020;12: 62.

79. Maunsell JHR. Neuronal representations of cognitive state: reward or attention? Trends Cogn Sci. 2004;8: 261–265.

80. Semedo JD, Zandvakili A, Machens CK, Yu BM, Kohn A. Cortical Areas Interact through a Communication Subspace. Neuron. 2019;102: 249–259.e4.

81. Huk AC. Multiplexing in the primate motion pathway. Vision Research. 2012. pp. 173–180. doi:10.1016/j.visres.2012.04.007

82. McIntosh AR, Mišić B. Multivariate statistical analyses for neuroimaging data. Annu Rev Psychol. 2013;64: 499–525.

83. Smith SM, Nichols TE, Vidaurre D, Winkler AM, Behrens TEJ, Glasser MF, et al. A positive-negative mode of population covariation links brain connectivity, demographics and behavior. Nat Neurosci. 2015;18: 1565–1567.

84. Llera A, Wolfers T, Mulders P, Beckmann CF. Inter-individual differences in human brain structure and morphology link to variation in demographics and behavior. Elife. 2019;8. doi:10.7554/eLife.44443

85. Kong TS, Gratton C, Low KA, Tan CH, Chiarelli AM, Fletcher MA, et al. Age-related differences in functional brain network segregation are consistent with a cascade of cerebrovascular, structural, and cognitive effects. Netw Neurosci. 2020;4: 89–114.

86. Morgan SE, White SR, Bullmore ET, Vértes PE. A Network Neuroscience Approach to Typical and Atypical Brain Development. Biol Psychiatry Cogn Neurosci Neuroimaging. 2018;3: 754–766.

87. Davis SW, Szymanski A, Boms H, Fink T, Cabeza R. Cooperative contributions of structural and functional connectivity to successful memory in aging. Netw Neurosci. 2019;3: 173–194.

88. Daianu M, Jahanshad N, Nir TM, Toga AW, Jack CR Jr, Weiner MW, et al. Breakdown of brain connectivity between normal aging and Alzheimer’s disease: a structural k-core network analysis. Brain Connect. 2013;3: 407–422.

89. Avesani P, McPherson B, Hayashi S, Caiafa C, Henschel R, Garyfallidis E, et al. The open diffusion data derivatives, brain data upcycling via integrated publishing of derivatives and reproducible open cloud services. Nature Scientific Data. 2019;6: 1–13.

90. Shafto MA, Tyler LK, Dixon M, Taylor JR, Rowe JB, Cusack R, et al. The Cambridge Centre for Ageing and Neuroscience (Cam-CAN) study protocol: a cross-sectional, lifespan, multidisciplinary examination of healthy cognitive ageing. BMC Neurol. 2014;14: 204.

91. Rubinov M, Sporns O. Complex network measures of brain connectivity: uses and interpretations. Neuroimage. 2010;52: 1059–1069.

92. Diedrichsen J, Kriegeskorte N. Representational models: A common framework for understanding encoding, pattern-component, and representational-similarity analysis. PLoS Comput Biol. 2017;13: e1005508.

93. Yeo BTT, Krienen FM, Sepulcre J, Sabuncu MR, Lashkari D, Hollinshead M, et al. The organization of the human cerebral cortex estimated by intrinsic functional connectivity. J Neurophysiol. 2011;106: 1125–1165.

94. Glasser MF, Coalson TS, Robinson EC, Hacker CD, Harwell J, Yacoub E, et al. A multi-modal parcellation of human cerebral cortex. Nature. 2016;536: 171–178.

95. Patenaude B, Smith SM, Kennedy DN, Jenkinson M. A Bayesian model of shape and appearance for subcortical brain segmentation. NeuroImage. 2011. pp. 907–922. doi:10.1016/j.neuroimage.2011.02.046

96. Ades-Aron B, Veraart J, Kochunov P, McGuire S, Sherman P, Kellner E, et al. Evaluation of the accuracy and precision of the diffusion parameter EStImation with Gibbs and NoisE removal pipeline. Neuroimage. 2018;183: 532–543.

97. Takemura H, Caiafa CF, Wandell BA, Pestilli F. Ensemble Tractography. PLoS Comput Biol. 2016;12: e1004692.

98. Smith RE, Tournier J-D, Calamante F, Connelly A. Anatomically-constrained tractography: Improved diffusion MRI streamlines tractography through effective use of anatomical information. NeuroImage. 2012. pp. 1924–1938. doi:10.1016/j.neuroimage.2012.06.005

99. Roberts JA, Perry A, Roberts G, Mitchell PB, Breakspear M. Consistency-based thresholding of the human connectome. NeuroImage. 2017. pp. 118–129. doi:10.1016/j.neuroimage.2016.09.053

100. Cheng H, Wang Y, Sheng J, Sporns O, Kronenberger WG, Mathews VP, et al. Optimization of seed density in DTI tractography for structural networks. J Neurosci Methods. 2012;203: 264–272.

101. Folstein MF. The Mini-Mental State Examination. Archives of General Psychiatry. 1983. p. 812. doi:10.1001/archpsyc.1983.01790060110016

102. Folstein MF, Folstein SE, McHugh PR. Mini-Mental State Examination. PsycTESTS Dataset. 2014. doi:10.1037/t07757-000

103. van den Heuvel MP, Sporns O. Rich-club organization of the human connectome. J Neurosci. 2011;31: 15775–15786.

104. Baggio HC, Segura B, Junque C, de Reus MA, Sala-Llonch R, Van den Heuvel MP. Rich Club Organization and Cognitive Performance in Healthy Older Participants. J Cogn Neurosci. 2015;27: 1801–1810.

105. Kriegeskorte N. Representational similarity analysis – connecting the branches of systems neuroscience. Frontiers in Systems Neuroscience. 2008. doi:10.3389/neuro.06.004.2008

106. Hao X, Li C, Yan J, Yao X, Risacher SL, Saykin AJ, et al. Identification of associations between genotypes and longitudinal phenotypes via temporally-constrained group sparse canonical correlation analysis. Bioinformatics. 2017;33: i341–i349.

107. Rex DE, Ma JQ, Toga AW. The LONI Pipeline Processing Environment. Neuroimage. 2003;19: 1033–1048.

108. Mueller SG, Weiner MW, Thal LJ, Petersen RC, Jack CR, Jagust W, et al. Ways toward an early diagnosis in Alzheimer’s disease: The Alzheimer’s Disease Neuroimaging Initiative (ADNI). Alzheimer’s & Dementia. 2005. pp. 55–66. doi:10.1016/j.jalz.2005.06.003

109. Weiner MW, Veitch DP, Aisen PS, Beckett LA, Cairns NJ, Green RC, et al. Recent publications from the Alzheimer’s Disease Neuroimaging Initiative: Reviewing progress toward improved AD clinical trials. Alzheimer’s & Dementia. 2017. pp. e1–e85. doi:10.1016/j.jalz.2016.11.007

110. Van Essen DC, Ugurbil K. The future of the human connectome. NeuroImage. 2012. pp. 1299–1310. doi:10.1016/j.neuroimage.2012.01.032

111. Thompson PM, Andreassen OA, Arias-Vasquez A, Bearden CE, Boedhoe PS, Brouwer RM, et al. ENIGMA and the individual: Predicting factors that affect the brain in 35 countries worldwide. Neuroimage. 2017;145: 389–408.

112. Shirer WR, Ryali S, Rykhlevskaia E, Menon V, Greicius MD. Decoding subject-driven cognitive states with whole-brain connectivity patterns. Cereb Cortex. 2012;22: 158–165.

113. Beckmann CF, Smith SM. Probabilistic Independent Component Analysis for Functional Magnetic Resonance Imaging. IEEE Transactions on Medical Imaging. 2004. pp. 137–152. doi:10.1109/tmi.2003.822821

114. Dahne S, Bieszmann F, Samek W, Haufe S, Goltz D, Gundlach C, et al. Multivariate Machine Learning Methods for Fusing Multimodal Functional Neuroimaging Data. Proceedings of the IEEE. 2015. pp. 1507–1530. doi:10.1109/jproc.2015.2425807

115. Silva RF, Plis SM, Adalı T, Calhoun VD. A statistically motivated framework for simulation of stochastic data fusion models applied to multimodal neuroimaging. Neuroimage. 2014;102 Pt 1: 92–117.

116. Lydon-Staley DM, Cornblath EJ, Blevins AS, Bassett DS. Modeling brain, symptom, and behavior in the winds of change. Neuropsychopharmacology. 2020. doi:10.1038/s41386-020-00805-6

117. Calhoun VD, Liu J, Adali T. A review of group ICA for fMRI data and ICA for joint inference of imaging, genetic, and ERP data. Neuroimage. 2009;45: S163–72.

118. Allen EA, Erhardt EB, Damaraju E, Gruner W, Segall JM, Silva RF, et al. A baseline for the multivariate comparison of resting-state networks. Front Syst Neurosci. 2011;5: 2.

119. Calhoun VD, Adali T, Pearlson GD, Pekar JJ. A method for making group inferences from functional MRI data using independent component analysis. Hum Brain Mapp. 2001;14: 140–151.

120. Du L, Liu F, Liu K, Yao X, Risacher SL, Han J, et al. Identifying diagnosis-specific genotype–phenotype associations via joint multitask sparse canonical correlation analysis and classification. Bioinformatics. 2020. pp. i371–i379. doi:10.1093/bioinformatics/btaa434

121. Hawrylycz MJ, Lein ES, Guillozet-Bongaarts AL, Shen EH, Ng L, Miller JA, et al. An anatomically comprehensive atlas of the adult human brain transcriptome. Nature. 2012;489: 391–399.

122. Krishnan A, Williams LJ, McIntosh AR, Abdi H. Partial Least Squares (PLS) methods for neuroimaging: a tutorial and review. Neuroimage. 2011;56: 455–475.

123. McIntosh AR, Lobaugh NJ. Partial least squares analysis of neuroimaging data: applications and advances. Neuroimage. 2004;23 Suppl 1: S250–63.

124. Bijsterbosch JD, Beckmann CF, Woolrich MW, Smith SM, Harrison SJ. The relationship between spatial configuration and functional connectivity of brain regions revisited. Elife. 2019;8. doi:10.7554/eLife.44890

125. Bae E, Hur J-W, Kim J, Kwon JS, Lee J, Lee S-H, et al. Multi-group analysis using generalized additive kernel canonical correlation analysis. Sci Rep. 2020;10: 12624.

126. Wang H-T, Smallwood J, Mourao-Miranda J, Xia CH, Satterthwaite TD, Bassett DS, et al. Finding the needle in a high-dimensional haystack: Canonical correlation analysis for neuroscientists. NeuroImage. 2020. p. 116745. doi:10.1016/j.neuroimage.2020.116745

127. Marek S, Tervo-Clemmens B, Calabro FJ, Montez DF, Kay BP, Hatoum AS, et al. Towards Reproducible Brain-Wide Association Studies. 2020. p. 2020.08.21.257758. doi:10.1101/2020.08.21.257758

128. Helmer M, Warrington S, Mohammadi-Nejad A-R, Ji JL, Howell A, Rosand B, et al. On stability of Canonical Correlation Analysis and Partial Least Squares with application to brain-behavior associations. 2020. p. 2020.08.25.265546. doi:10.1101/2020.08.25.265546

129. Mueller RO, Hancock GR. Structural Equation Modeling. The Reviewer’s Guide to Quantitative Methods in the Social Sciences. 2018. pp. 445–456. doi:10.4324/9781315755649-33

130. Schulz M-A, Thomas Yeo BT, Vogelstein JT, Mourao-Miranada J, Kather JN, Kording K, et al. Different scaling of linear models and deep learning in UKBiobank brain images versus machine-learning datasets. Nature Communications. 2020. doi:10.1038/s41467-020-18037-z

131. Assem M, Blank IA, Mineroff Z, Ademoğlu A, Fedorenko E. Activity in the fronto-parietal multiple-demand network is robustly associated with individual differences in working memory and fluid intelligence. Cortex. 2020;131: 1–16.

132. Hsu LL, Culhane AC. Impact of Data Preprocessing on Integrative Matrix Factorization of Single Cell Data. Front Oncol. 2020;10: 973.

133. Fischl B. FreeSurfer. NeuroImage. 2012. pp. 774–781. doi:10.1016/j.neuroimage.2012.01.021

134. Tournier J-D, Smith R, Raffelt D, Tabbara R, Dhollander T, Pietsch M, et al. MRtrix3: A fast, flexible and open software framework for medical image processing and visualisation. Neuroimage. 2019;202: 116137.

135. Greve DN, Fischl B. Accurate and robust brain image alignment using boundary-based registration. Neuroimage. 2009;48: 63–72.

136. Jeurissen B, Tournier J-D, Dhollander T, Connelly A, Sijbers J. Multi-tissue constrained spherical deconvolution for improved analysis of multi-shell diffusion MRI data. NeuroImage. 2014. pp. 411–426. doi:10.1016/j.neuroimage.2014.07.061

137. Anderson C. Docker [Software engineering]. IEEE Software. 2015. pp. 102–c3. doi:10.1109/ms.2015.62

138. Kurtzer GM, Sochat V, Bauer MW. Singularity: Scientific containers for mobility of compute. PLoS One. 2017;12: e0177459.

139. Guntupalli JS, Hanke M, Halchenko YO, Connolly AC, Ramadge PJ, Haxby JV. A Model of Representational Spaces in Human Cortex. Cereb Cortex. 2016;26: 2919–2934.

