## Supplemental Figures for "A single-mode associates global patterns of brain network structure and behavior across the human lifespan"

### Author Information

1. Department of Psychological and Brain Sciences, Indiana University Bloomington, 1101 E 10th Street, Bloomington, Indiana USA 47405.
2. Department of Psychology, The University of Texas at Austin, 108 E. Dean Keeton St., Austin, TX 78712

### Abstract

Multiple human behaviors improve early in life, peaking in young adulthood, and declining thereafter. Several properties of brain structure and function progress similarly across the lifespan. Cognitive and neuroscience research has approached aging primarily using associations between a few behaviors, brain functions, and structures. Because of this, the multivariate, global factors relating brain and behavior across the lifespan are not well understood. We investigated the global patterns of associations between 334 behavioral and clinical measures and 376 brain structural connections in 594 individuals across the lifespan. A single-axis associated changes in multiple behavioral domains and brain structural connections (r=0.5808). Individual variability within the single association axis well predicted the age of the subject (r=0.6275). Representational similarity analysis evidenced global patterns of interactions across multiple brain network systems and behavioral domains. Results show that global processes of human aging can be well captured by a multivariate data fusion approach. [147]

#### Data availability

The source data are provided by the Cambridge Aging Neuroscience Project https://camcan-archive.mrc-cbu.cam.ac.uk/. Brain data derived as part of this project and used as features for all the analyses are available on brainlife.io/pubs: <DOI>

#### Code availability

Code is available on github at https://github.com/bcmcpher/cca_aging and as web services reproducing the analyes at <URL>

#### Acknowledgments

This research was supported by NSF OAC-1916518, NSF IIS-1912270, NSF IIS-1636893, NSF BCS-1734853, Microsoft Faculty Fellowship to F.P., NIH 5T32MH103213-05 to William Hetrick. We thank Soichi Hayashi, and Josh Faskowitz for contributing to the development of brainlife.io, Craig Stewart, Robert Henschel, David Hancock and Jeremy Fischer for support with jetstream-cloud.org (NSF ACI-1445604).

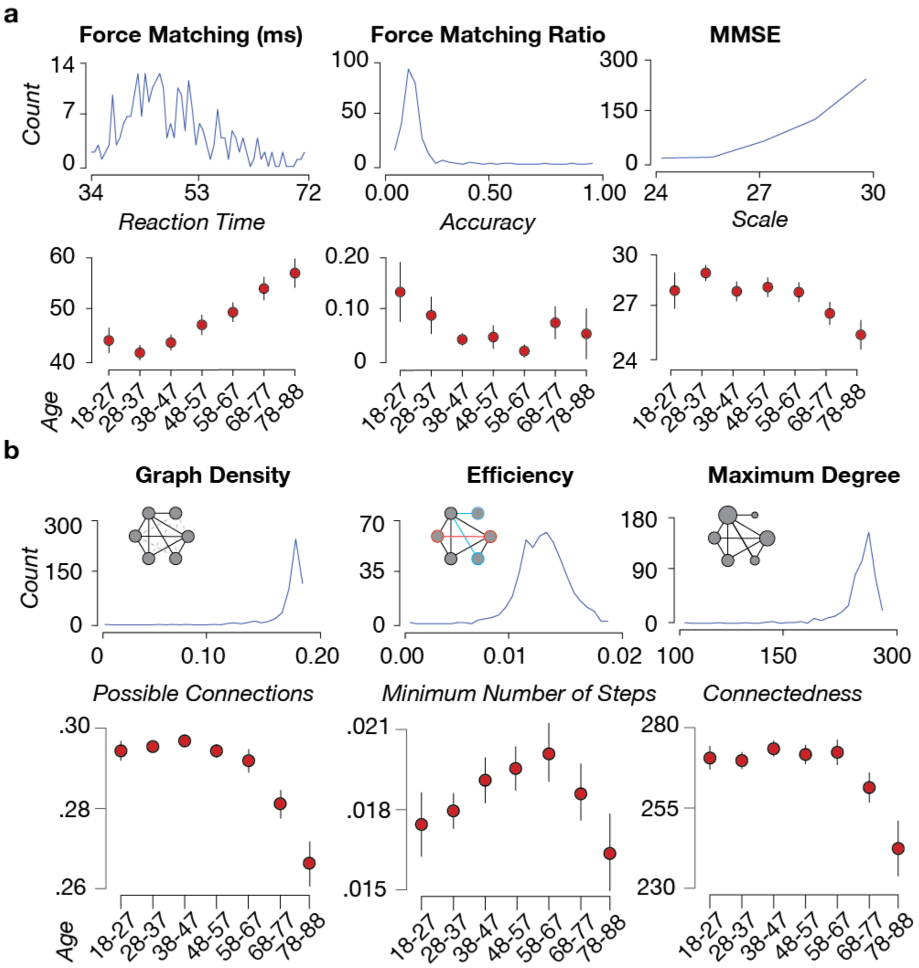


**Supplemental Figure 1. Variation across age of behavioral and network measures. a. Behavior histograms and binned averages across age.** Example histogram of the reaction times in the force matching task (*top left;* Shafto et al., 2014), the accuracy of matching the force (*top center;* Shafto et al., 2014), and the mini-mental state exam (MMSE; Shafto, et al., 2014; *top right*) are shown from the sample. Average and ±1 s.d. for each behavioral task, respectively. Data was binned in decades of subjects' age (*bottom row*). **b. Network histograms and binned averages across age.** Example histogram of the graph density (*bottom left*), the graph efficiency (*bottom center*), and the maximum node degree (*bottom right*) are shown from the sample. The average values and variance are binned for each decade, showing the mean and 2 units of standard error for the respective measure (*bottom row*).

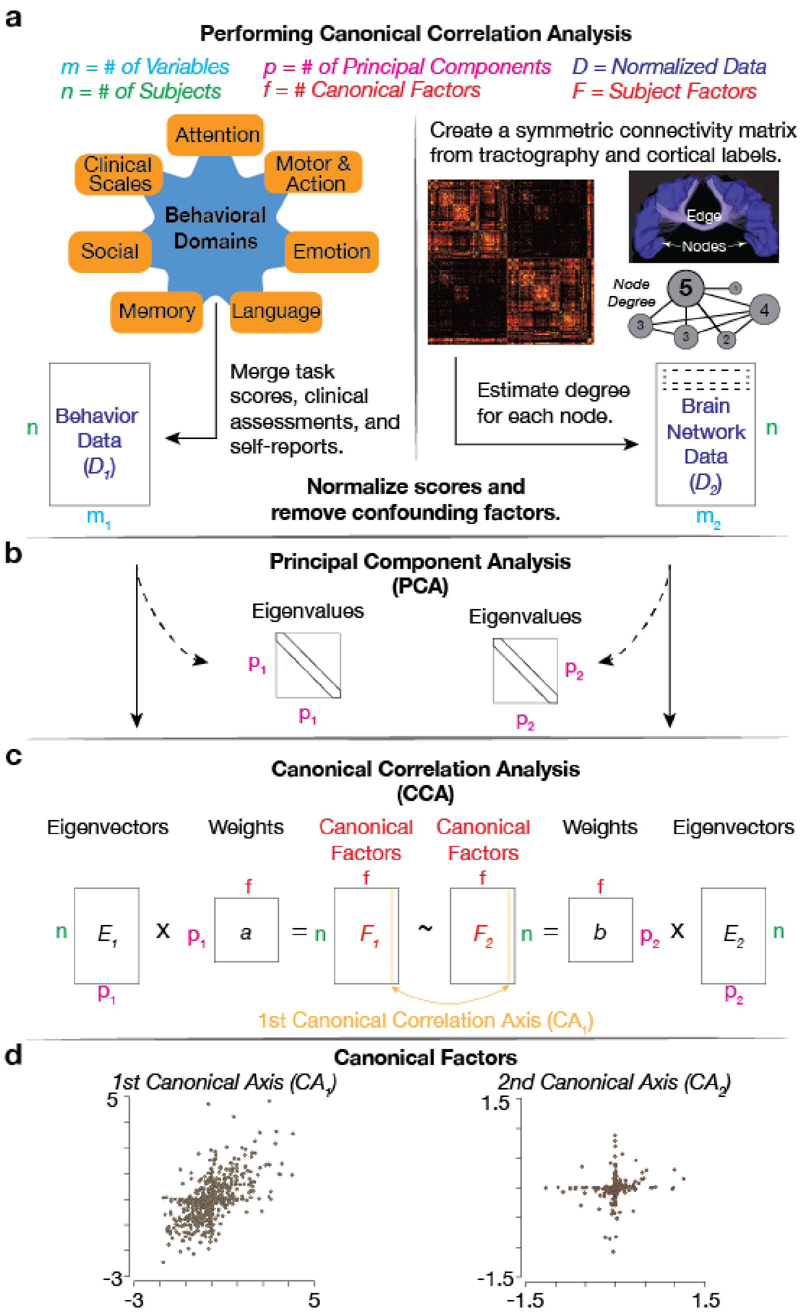


**Supplemental Figure 2a, b, c, and d. The flow of data through the analysis. a. Behavioral domains for tasks and questionnaires.** Variables estimates by the CAN consortium from multiple behavioral domains (*left*, *blue and orange*) were collected for each subject and organized into a matrix ($D_1$) with 594 ($n$) subjects and 334 variables ($m_1$). Node degree was estimated for each subject's brain network matrix (*right, black and orange*). An example pair of nodes and an edge is shown, along with a ball and stick diagram showing the values of node degree. The measures of node degree are a vector of 376 entries per subject. The node degree vectors for each subject were stacked to build the Brain Network Data matrix ($D_2$). $D_1$ and $D_2$ were normalized by computing the z-scores by columns. **b. Principal component analysis for dimensionality reduction.** The matrices $D_1$ and $D_2$ have size 594 X 334 and 594 X 376, respectively. A principal component analysis was performed independently for $D_1$ and $D_2$ to reduce the large sets of variables into a smaller set of components that still predicted most of the variance in the data (Smith et al., 2015). We estimated Eigenvalues and Eigenvectors from $D_1$ and $D_2$, and used the eigenvector matrices as data for a canonical correlation analysis (CCA, see next). We note that we performed multiple PCAs, with different numbers of principal components and used the number of principal components to tune the model prediction of the subjects age (See **Supplemental Figure 2e-g**). **c. Performing the canonical correlation.** The eigenvectors matrices obtained via PCA ($E_1$ and $E_2$) were used as input data to a CCA analysis. CCA estimated the inner weights ($a$ and $b$) and canonical factors ($F_1$ and $F_2$) simultaneously using the behavioral and brain network eigenvectors that maximized the correlation between the two input matrices ($E_1$ and $E_2$). The correlation from the first component, second and subsequent components estimated in $F_1$ and $F_2$ and the focus of analysis (See **Figure 2** and **3** and associated Supplemental Figures). **d. Plotting the CCA axis.** Example of first canonical axis (light orange) and second canonical axis (dark orange) estimated from the CCA.

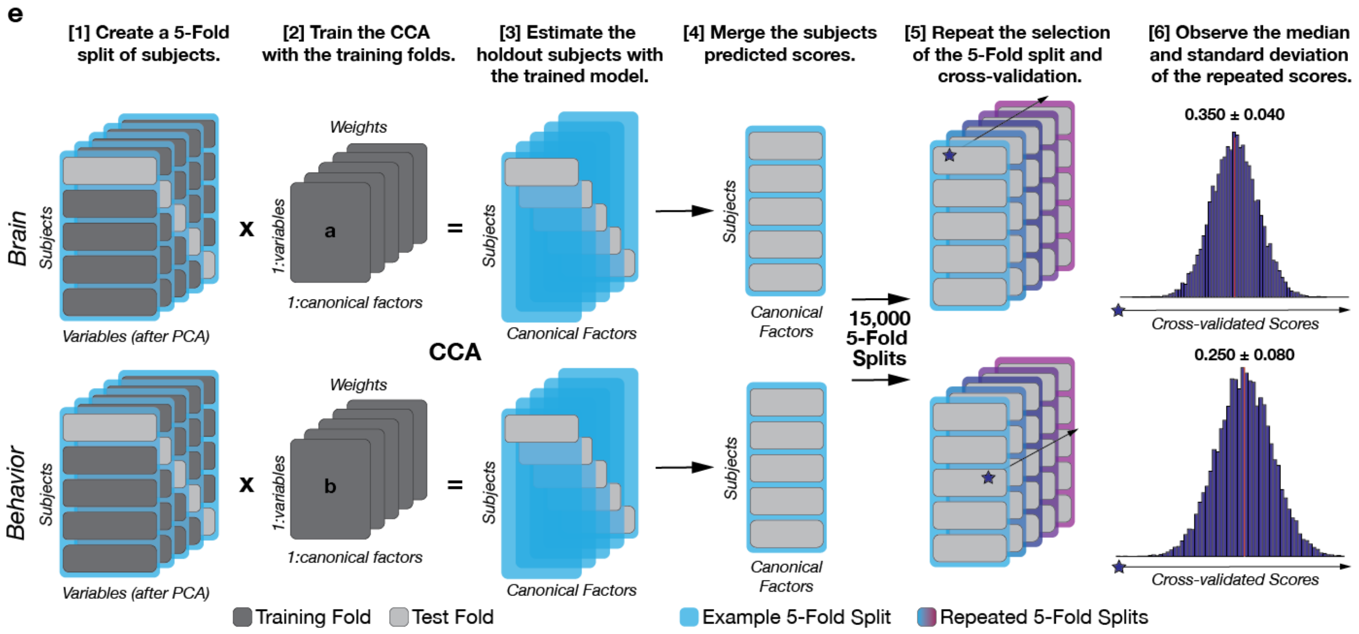


**Supplemental Figure 2e. Approach to repeated 5-fold cross-validation for the canonical correlation analysis (CCA).** We used 5-fold cross-validation to find the CCA model that simultaneously best predicted the brain networks and behavioral data. To do so, we split the data into 5 groups of subjects (*[1] dark and light gray*). Eighty percent of the data (*[1] dark gray*) was used to estimate the inner weights of the CCA model for both the behavioral and the brain network data – *[2]* **a** and **b** (see also **Supplementary Figure 2c**). The estimated weights were then used to estimate the CCA factors for the remaining, left-out 20% of the data (*[3] cyan and light grey*). The canonical factors for the complete set of subjects were estimated by repeating this process five times shifting the subjects utilized during each cross-validation fold (compare [1], [2], [3] and [4]). The predicted scores for each subject were then combined into a single estimate of all the predicted factors for every subject *[4]*. This 5-fold cross-validation process *[1-4]* was repeated 15,000, each time utilizing different 80/20 splits of the subjects. This resulted in 15,000 estimates of each canonical factor *[5]*. Finally, the median (*red line*) and standard deviation of the estimated canonical factors were computed for each distribution of 15,000 estimates [6]. Blue stars represent the example factor used to build the distributions.

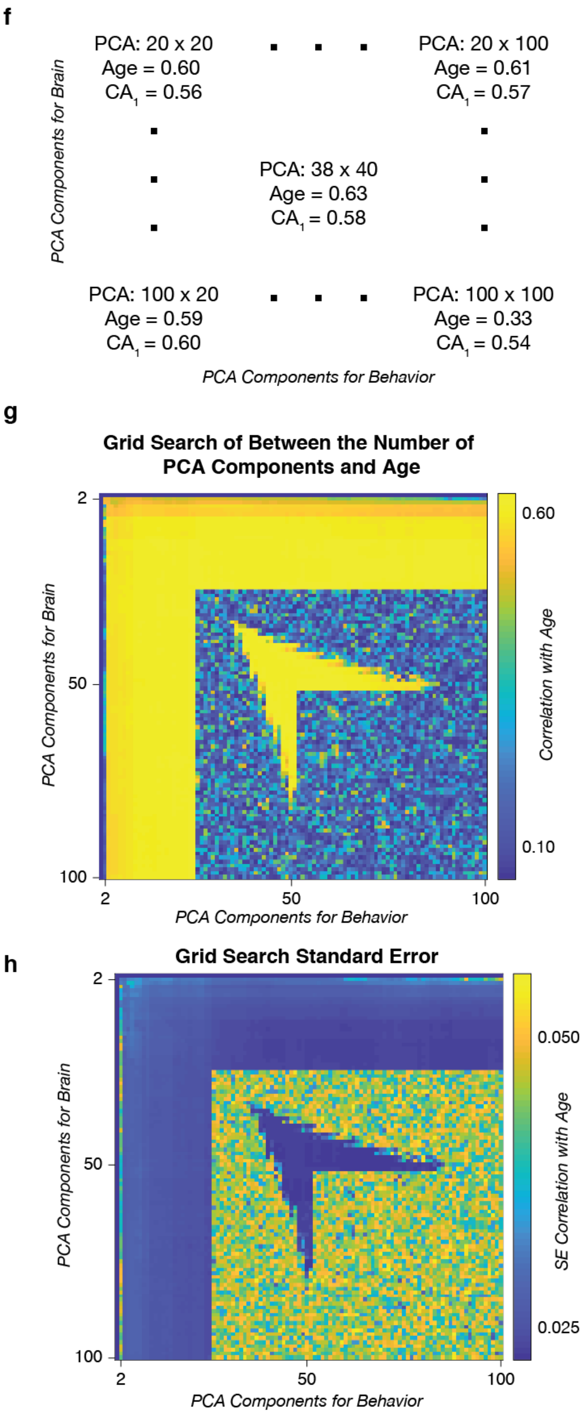


**Supplemental Figure 2f, g** and **h. CCA Model selection via cross-validated grid search of PCs.**

**f. A conceptual diagram of the PCA tuning space.** A simplified diagram illustrates the grid search that was performed to test different numbers of principal components. Every pair of PCA components between 2 and 100 was estimated during the processing of the data. The resulting canonical correlation and the correlation of the canonical axis with age are reported. The highest correlation, at 38 x 40 was selected for interpretation.

**g. The entire search space of the parameter tuning.** This shows the correlation of age for every parameter combination tested in the grid search. Yellow values indicate a high correlation with age while blue indicates the correlation between the datasets is low. We selected the PCA pair with the highest correlation.

**h. The standard error estimate of the correlation.** The standard error estimate of the correlation with age. The darker blue the value, the lower the standard error and the more confidence in the estimate.

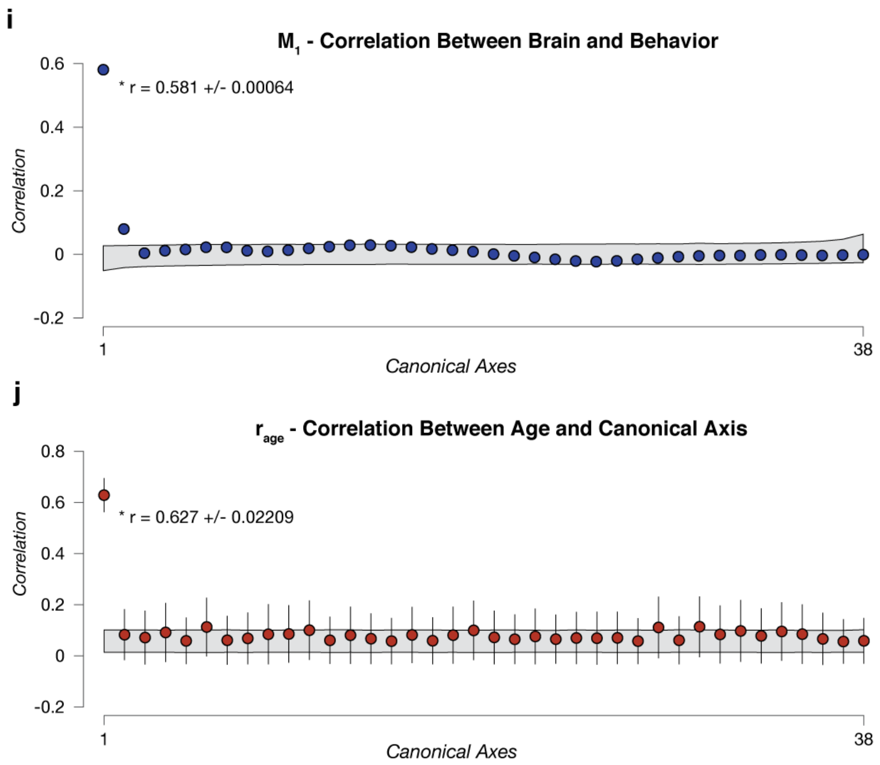


**Supplemental Figure 2i and j. All canonical correlations and all axis correlations with age for $M_1$.**

**i. All correlations estimated for each canonical axis for the final tuned PCA selection.** Each circle (*blue*) represents the cross-validated correlation estimated along the canonical axis. Each error bar represents 2 units of standard error for the estimate obtained via cross-validation. The gray band represents the 5th and 95th percentiles of a bootstrapped null distribution.

**j. Correlation with age for each canonical axis for the final tuned PCA selection.** Each circle (*red*) represents the correlation of age to the estimated canonical axis of each canonical factor. Each error bar represents 2 units of standard error for the estimate obtained via cross-validation. The gray band represents the 5th and 95th percentiles of a bootstrapped null distribution.

k

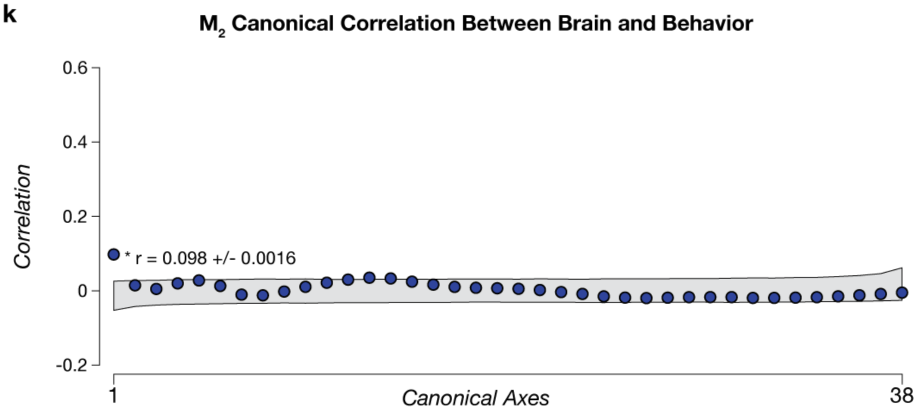


l

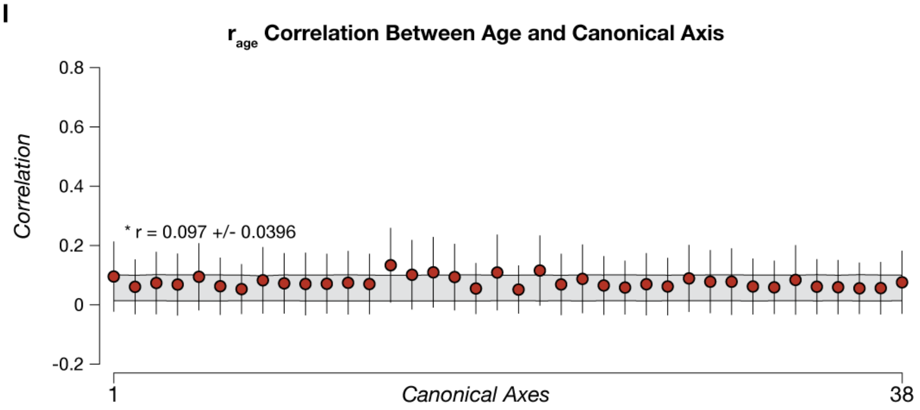


**Supplemental Figure 2k and l. All canonical correlations and all axis correlations with age for $M_2$.**

**k. All correlations estimated for each canonical axis for the final tuned PCA selection.** Each circle (*blue*) represents the cross-validated correlation estimated along the canonical axis. Each error bar represents 2 units of standard error for the estimate obtained via cross-validation. The gray band represents the 5th and 95th percentiles of a bootstrapped null distribution.

**l. Correlation with age for each canonical axis for the final tuned PCA selection.** Each circle (*red*) represents the correlation of age to the estimated canonical axis of each canonical factor. Each error bar represents 2 units of standard error for the estimate obtained via cross-validation. The gray band represents the 5th and 95th percentiles of a bootstrapped null distribution.

### Components of brain and behavior contributing to the CCA results.

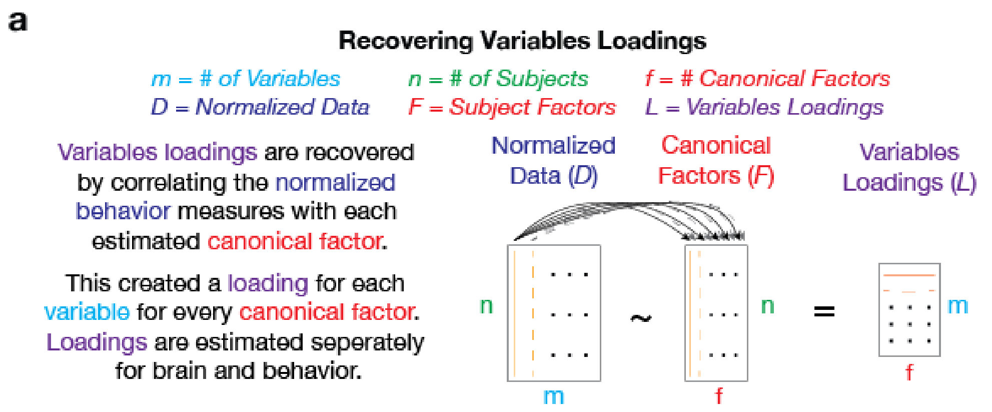


**Supplemental Figure 3a. The recovery of variables loadings.** To interpret how individual variables contribute to the CCA axis (*L, purple*), a correlation is taken between the original variables (*D, blue*) and the estimated canonical factors (*F, red*). By finding the correlation between each variable in *D* to a single factor in *F*, the loadings for every variable to the factor are recovered. By finding the correlation between a variable in *D* to every factor in *F*, the loading of the variables onto every factor is recovered. All variables by factor loadings were organized into matrix (*L*). These steps are performed for the brain and behavioral domain data ($D_1$ and $D_2$) independently to generate two variable loading matrices, $L_1$ and $L_2$.

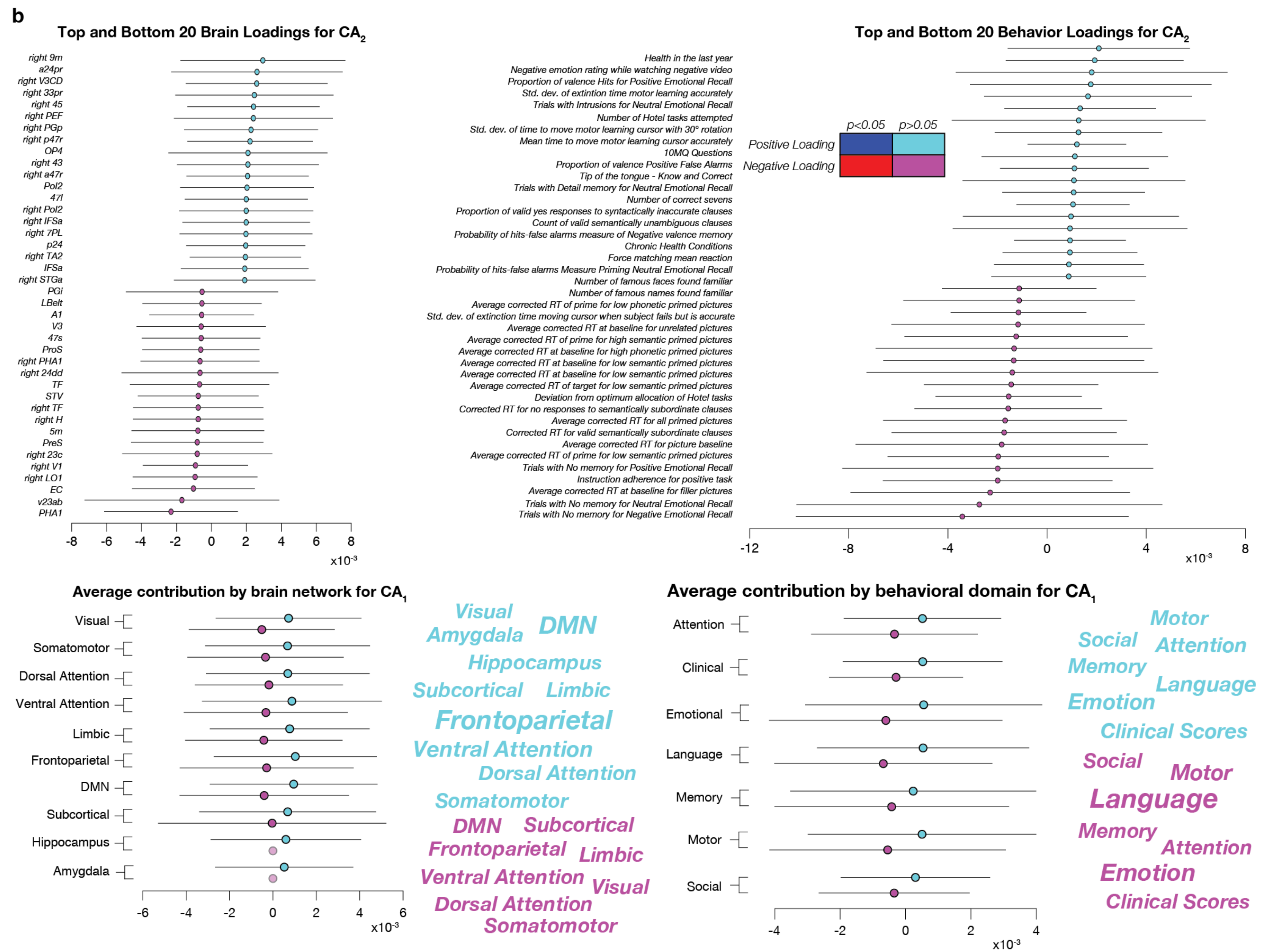
b
Top and Bottom 20 Brain Loadings for $CA_2$
right 9m
a24pr
right V3CD
right 33pr
right 45
right PEF
right PGp
right p47r
OP4
right 43
right a47r
Pol2
47l
right Pol2
right IFSa
right 7PL
p24
right TA2
IFSa
right STGa
PGi
LBelt
A1
V3
47s
ProS
right PHA1
right 24dd
TF
STV
right TF
right H
5m
PreS
right 23c
right V1
right LO1
EC
v23ab
PHA1
-8 -6 -4 -2 0 2 4 6 8
x10-3
Top and Bottom 20 Behavior Loadings for $CA_2$
Health in the last year
Negative emotion rating while watching negative video
Proportion of valence Hits for Positive Emotional Recall
Std. dev. of extintion time motor learning accurately
Trials with Intrusions for Neutral Emotional Recall
Number of Hotel tasks attempted
Std. dev. of time to move motor learning cursor with 30° rotation
Mean time to move motor learning cursor accurately
10MQ Questions
Proportion of valence Positive False Alarms
Tip of the tongue - Know and Correct
Trials with Detail memory for Neutral Emotional Recall
Number of correct sevens
Proportion of valid yes responses to syntactically inaccurate clauses
Count of valid semantically unambiguous clauses
Probability of hits-false alarms measure of Negative valence memory
Chronic Health Conditions
Force matching mean reaction
Probability of hits-false alarms Measure Priming Neutral Emotional Recall
Number of famous faces found familiar
Number of famous names found familiar
Average corrected RT of prime for low phonetic primed pictures
Std. dev. of extinction time moving cursor when subject fails but is accurate
Average corrected RT at baseline for unrelated pictures
Average corrected RT of prime for high semantic primed pictures
Average corrected RT at baseline for high phonetic primed pictures
Average corrected RT at baseline for low semantic primed pictures
Average corrected RT at baseline for low semantic primed pictures
Average corrected RT of target for low semantic primed pictures
Deviation from optimum allocation of Hotel tasks
Corrected RT for no responses to semantically subordinate clauses
Average corrected RT for all primed pictures
Corrected RT for valid semantically subordinate clauses
Average corrected RT for picture baseline
Average corrected RT of prime for low semantic primed pictures
Trials with No memory for Positive Emotional Recall
Instruction adherence for positive task
Average corrected RT at baseline for filler pictures
Trials with No memory for Neutral Emotional Recall
Trials with No memory for Negative Emotional Recall
p<0.05 p>0.05
Positive Loading
Negative Loading
-12 -8 -4 0 4 8
x10-3
Average contribution by brain network for $CA_1$
Visual
Somatomotor
Dorsal Attention
Ventral Attention
Limbic
Frontoparietal
DMN
Subcortical
Hippocampus
Amygdala
-6 -4 -2 0 2 4 6
x10-3
Visual DMN
Amygdala
Hippocampus
Subcortical Limbic
Frontoparietal
Ventral Attention
Dorsal Attention
Somatomotor
DMN Subcortical
Frontoparietal Limbic
Ventral Attention Visual
Dorsal Attention
Somatomotor
Average contribution by behavioral domain for $CA_1$
Attention
Clinical
Emotional
Language
Memory
Motor
Social
-4 -2 0 2 4
x10-3
Motor
Social Attention
Memory
Language
Emotion
Clinical Scores
Social Motor
Language
Memory
Attention
Emotion
Clinical Scores

**Supplemental Figure 3b. Loadings for the second canonical axis ($CA_2$) of $M_1$.** The arrows represent the strength of the correlation of the variable to the second canonical axis. Blue and red arrows represent positive and negative loadings significantly different from 0 while cyan and pink represent positive and negative loadings that are not significantly different from 0. Despite **$CA_2$** being marginally outside the range of the null distribution (**Supplemental Figure 2k**), there are no cross-validated loadings that are significantly different from zero that contributed to this finding. Due to the lack of significantly contributing variable loadings in **$CA_2$** we focused on the single factor solution for our presented findings.

c

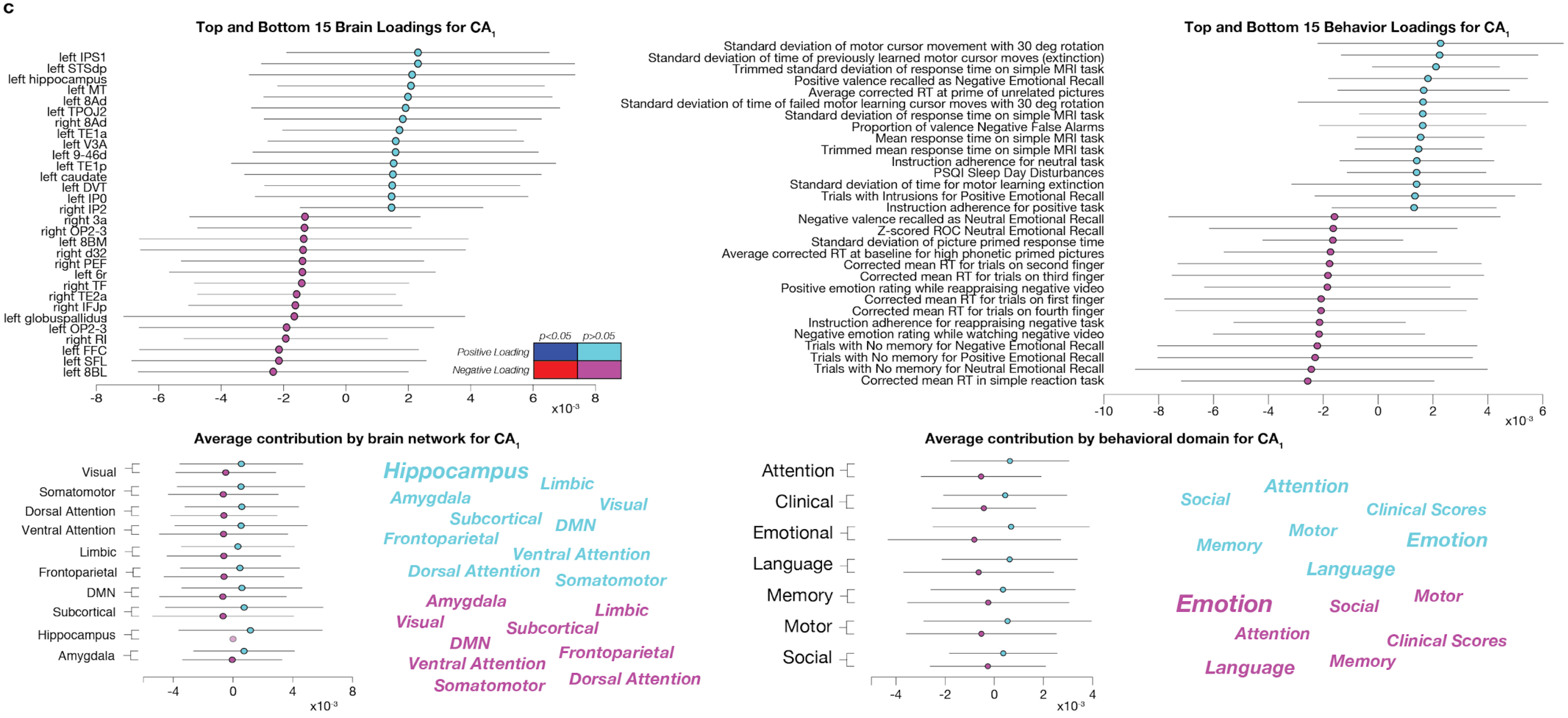

Top and Bottom 15 Brain Loadings for CA1
left IPS1
left STSdp
left hippocampus
left MT
left 8Ad
left TPOJ2
right 8Ad
left TE1a
left V3A
left 9-46d
left TE1p
left caudate
left DVT
left IP0
right IP2
right 3a
right OP2-3
left 8BM
right d32
right PEF
left 6r
right TF
right TE2a
right IFJp
left globuspallidus
left OP2-3
right RI
left FFC
left SFL
left 8BL
p<0.05
p>0.05
Positive Loading
Negative Loading
x10-3
Top and Bottom 15 Behavior Loadings for CA1
Standard deviation of motor cursor movement with 30 deg rotation
Standard deviation of time of previously learned motor cursor moves (extinction)
Trimmed standard deviation of response time on simple MRI task
Positive valence recalled as Negative Emotional Recall
Average corrected RT at prime of unrelated pictures
Standard deviation of time of failed motor learning cursor moves with 30 deg rotation
Standard deviation of response time on simple MRI task
Proportion of valence Negative False Alarms
Mean response time on simple MRI task
Trimmed mean response time on simple MRI task
Instruction adherence for neutral task
PSQI Sleep Day Disturbances
Standard deviation of time for motor learning extinction
Trials with Intrusions for Positive Emotional Recall
Instruction adherence for positive task
Negative valence recalled as Neutral Emotional Recall
Z-scored ROC Neutral Emotional Recall
Standard deviation of picture primed response time
Average corrected RT at baseline for high phonetic primed pictures
Corrected mean RT for trials on second finger
Corrected mean RT for trials on third finger
Positive emotion rating while reappraising negative video
Corrected mean RT for trials on first finger
Corrected mean RT for trials on fourth finger
Instruction adherence for reappraising negative task
Negative emotion rating while watching negative video
Trials with No memory for Negative Emotional Recall
Trials with No memory for Positive Emotional Recall
Trials with No memory for Neutral Emotional Recall
Corrected mean RT in simple reaction task
x10-3
Average contribution by brain network for CA1
Visual
Somatomotor
Dorsal Attention
Ventral Attention
Limbic
Frontoparietal
DMN
Subcortical
Hippocampus
Amygdala
x10-3
Hippocampus
Limbic
Amygdala
Visual
Subcortical
DMN
Frontoparietal
Ventral Attention
Dorsal Attention
Somatomotor
Amygdala
Limbic
Visual
Subcortical
DMN
Frontoparietal
Ventral Attention
Dorsal Attention
Somatomotor
Average contribution by behavioral domain for CA1
Attention
Clinical
Emotional
Language
Memory
Motor
Social
x10-3
Social
Attention
Clinical Scores
Memory
Motor
Emotion
Language
Emotion
Social
Motor
Attention
Clinical Scores
Language
Memory

**Supplemental Figure 3c. Loadings for the first canonical axis ($CA_1$) of $M_2$.** The arrows represent the strength of the correlation of the variable to the second canonical axis. Blue and red arrows represent positive and negative loadings significantly different from 0 while cyan and pink represent positive and negative loadings that are not significantly different from 0. Despite ***$CA_1$*** being marginally outside the range of the null distribution (**Supplemental Figure 2k**), there are no cross-validated loadings that are significantly different from zero that contributed to this finding. Due to the lack of significantly contributing variable loadings in **$CA_1$** we focused on the single factor solution for our presented findings.

The reader can compare the loadings for $M_1$ and $M_2$ by comparing **Figure 3a** and **b** and **Supplemental Figure 3b** and **c**. **Figure 3a** and **b** reports the loadings for $M_1$ $CA_1$. **Supplementary Figure 3b** reports the loadings for $M_1$ $CA_2$ and **Supplementary Figure 3c** reports the loadings for $M_2$ $CA_1$. See the table below for additional clarification.

| | ***$CA_1$*** | ***$CA_2$*** |
|---|---|---|
| ***$M_1$*** | Figure 3a and b | Supp Figure 3b |
| ***$M_2$*** | Supp Figure 3c | not reported (extremely small loadings) |

**Supplementary Table 1**. References to Figures containing Models and CCA axes.

The results show that whereas the weights for $M_1$ $CA_1$ are large and reliable, the weights for both $M_1$ $CA_2$ and $M_2$ $CA_1$ are much smaller and more variable. This comparison supports the hypothesis that a single axis predicting the quadratic trends across the lifespan (i.e., $M_1$ $CA_1$); hence when the CCA model is built without removing the participants' age. Opposite to that, if the participants' age is removed as in the case of M2 the CCA model fails in predicting a substantial portion of the variance in the data from the two domains.

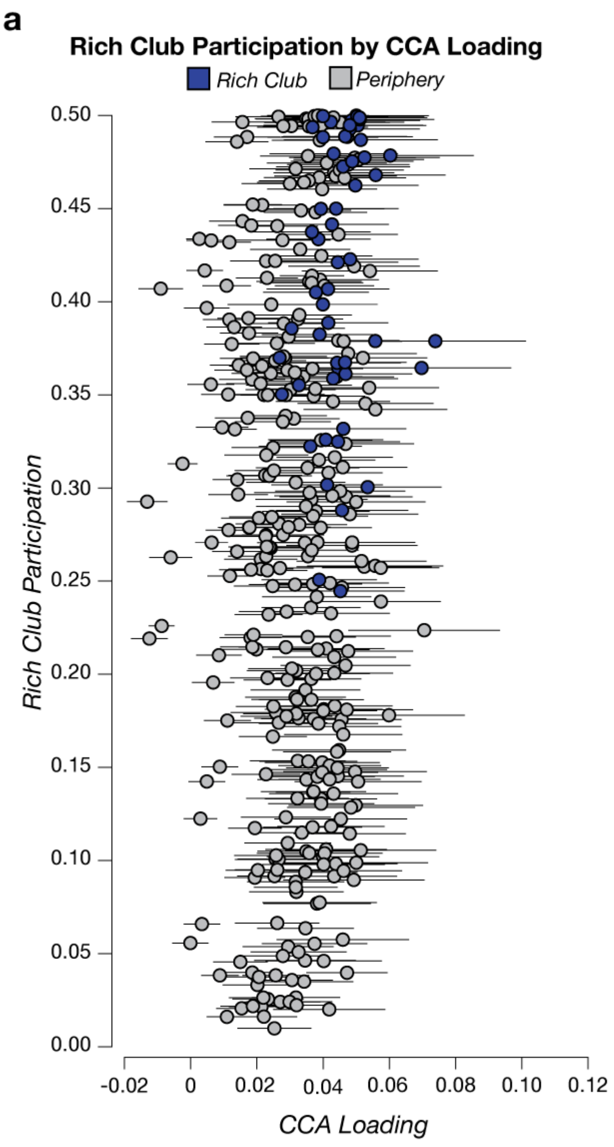


**Supplemental Figure 4a. The relationship of the rich club to the CCA loadings. a. Rich club participation compared to CCA loading.** Each symbol in the scatter plot represents a cortical region. Blue symbols are part of the core rich club. Gray symbols are part of the rich-club periphery. Error bars represent 2 s.e. estimated via cross-validation of the CCA loadings.

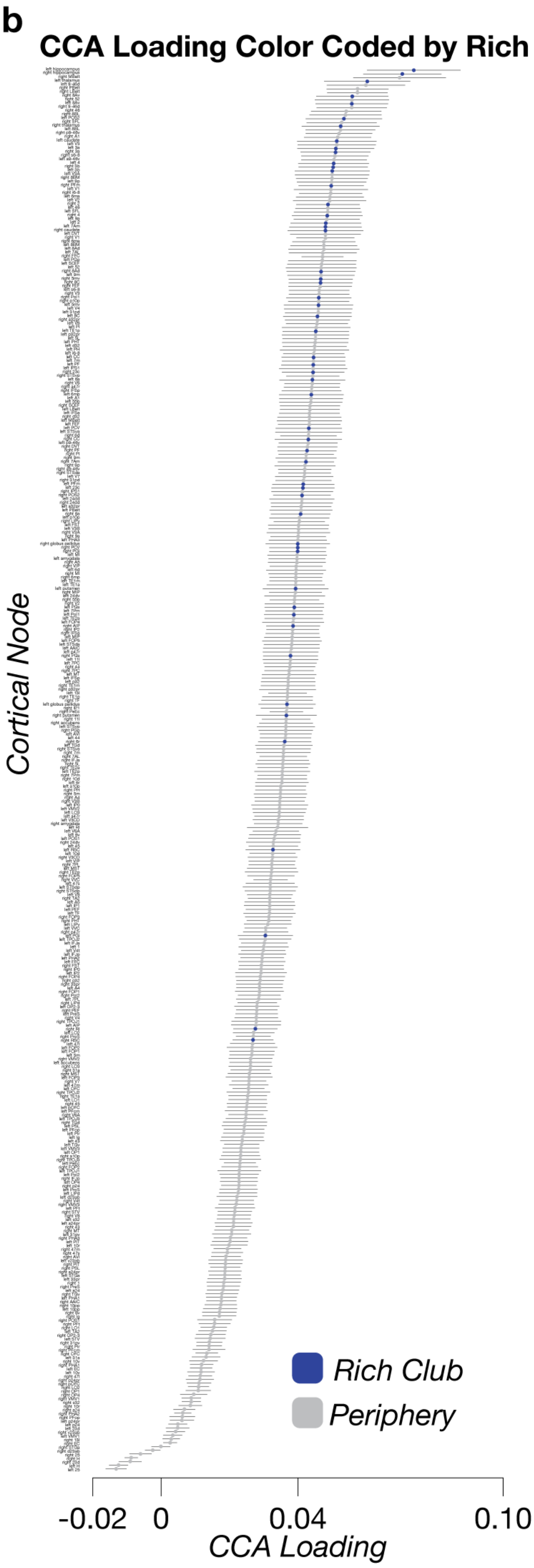


**Supplemental Figure 4b. Cortical regions sorted by CCA loading magnitude.**

Each symbol in the scatter plot represents a cortical region (mean $CA_1$ loading ±2 s.e.). The loadings for all regions were sorted from highest to lowest. Core rich club nodes are blue and periphery nodes are gray

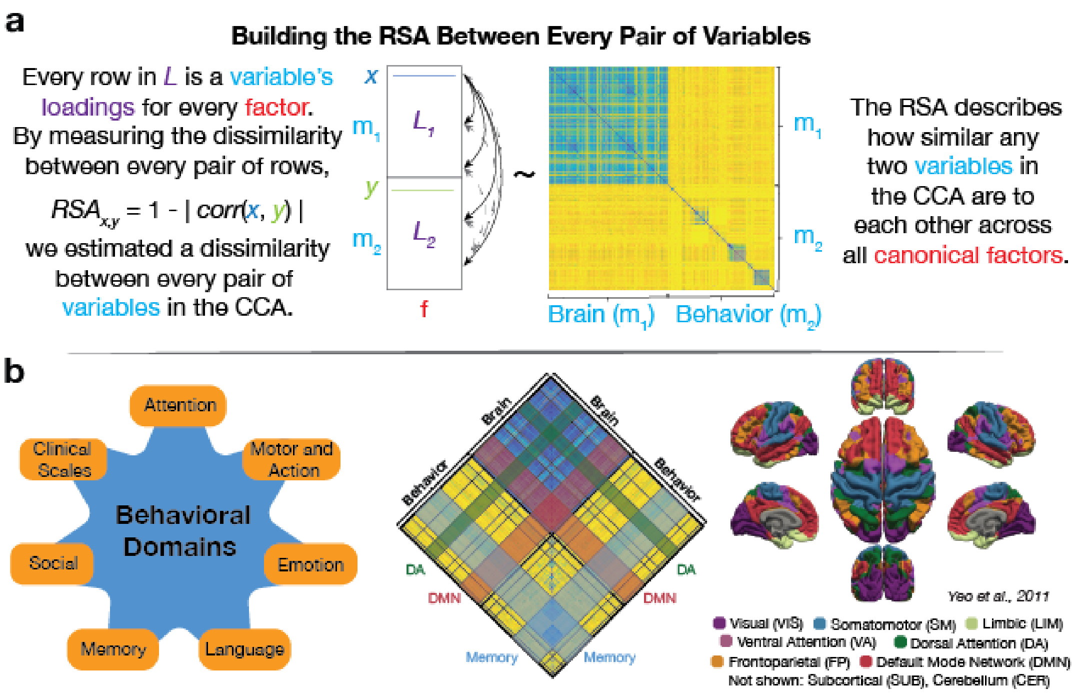


**Supplemental Figure 5a and b. Representational Similarity Analysis from Variables Loadings.**

**a. Estimating representational factor dissimilarity matrix ($S_1$).** To estimate the representational factor dissimilarity matrix ($S_1$) all variable loadings for each estimated canonical factor are required ($L_1$ and $L_2$). A dissimilarity between any pair of variables in the model (for example, *x* and *y*) is estimated by first computing the correlation (*r*) between the loadings of the variables across all the estimated factors (*f*) and computing *1 - |r|*, (Eq. 4). This value describes how dissimilar these variables are with each other across all estimated factors (*f*). By estimating these values for every pair of variables in the analysis, a full estimate of the dissimilarities is created (yellow and blue matrix).

**b. Grouping variables by domain.** We compute the mean of the RSA values within each behavioral domain (Shafto et al., 2014) and within major functional brain networks (Yeo et al. 2011). Behavioral domains are presented in orange around a blue star (from **Figure 1b,** *left*) and the nodes from the network parcellation were assigned to the Yeo atlas labels (*right*) based on the proportion overlap of the nodes in an atlas image. The dorsal attention network (DA, *green*), the default mode network (DMN, *red*), and the memory tasks (*blue*) are highlighted in the center dissimilarity matrix.

The RSA approach used here (**Supplemental Figure 5**) in turn allowed us to describe the simultaneous contributions of each variable to multiple other variables. The CCA variables loadings ($L_1$ and $L_2$) were used as inputs for a RSA. More specifically, every variable loading in $M_1$ (376 network and 334 behavior variables) was first correlated with the loadings of all the other variables. After that, the dissimilarity was then computed as *1-r*, see *Eq. 4*. This process generated a square, symmetric matrix, $S_1$ of size 710x710 (**Supplemental Figure 5c**, *top*).

Within this RSA framework, a low dissimilarity between the loadings of two variables would indicate that the variables contribute coherently to the CCA factors. Conversely, a high dissimilarity between two variable loadings would indicate that the variables contribute differently to the CCA. Because the CCA factors within each data domain (brain or behavior) are organized in descending order, the correlation within a single data domain is expected to be larger than that between data domains. $S_1$ allowed us to capture the coherence among the loadings of the variables and explore the cross-domain associations contributing to the CCA model. Furthermore, because $S_1$ was constructed using model $M_1$, in which the majority of the variance was explained by subject's age (see **Figure 2** and associated text), it was assumed that $S_1$ also depended on subject age.

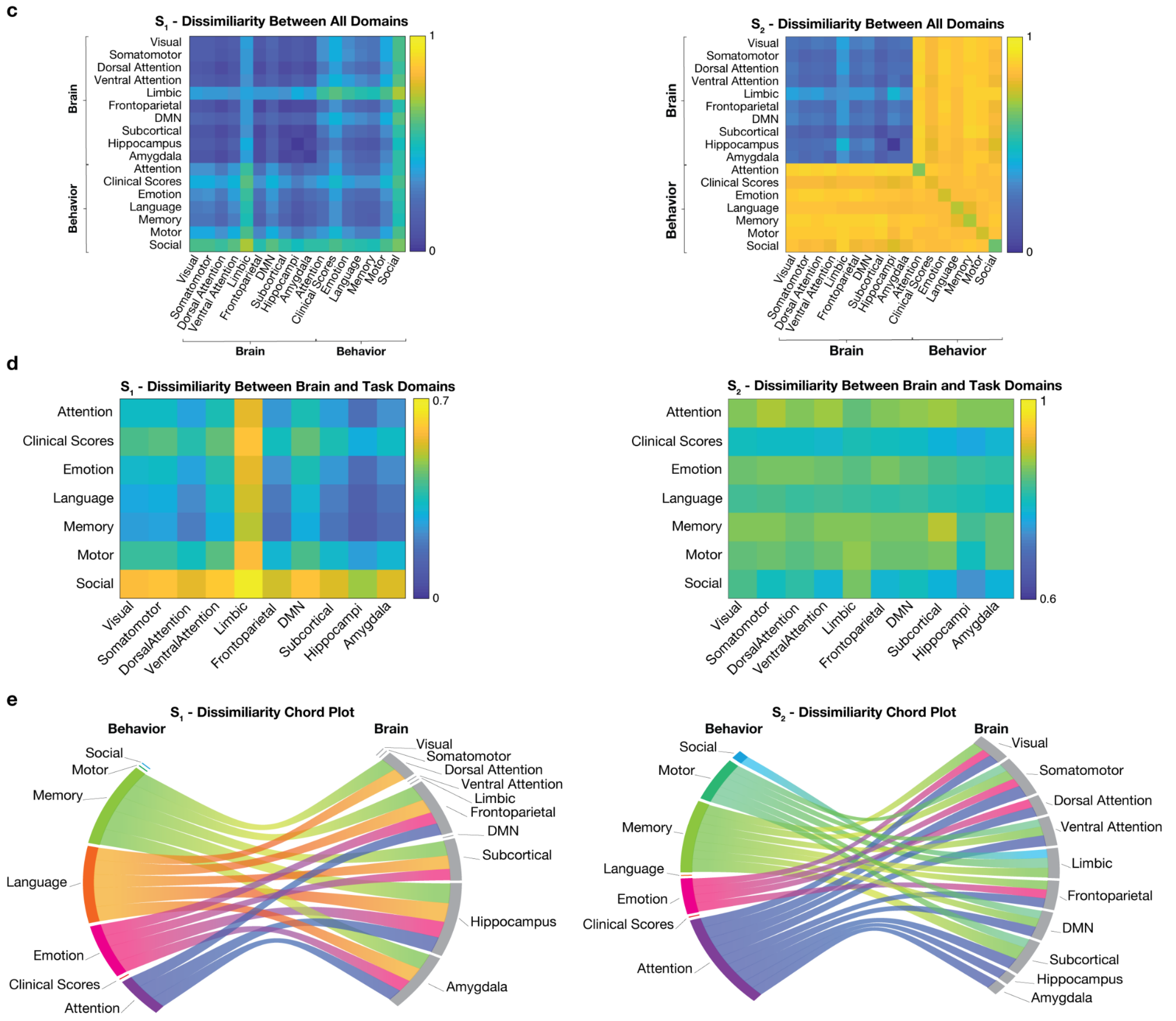


**Supplemental Figure 5c, d, and e. Comparing the difference between RSA with age not accounted for as a covariate and age accounted for as a covariate.**

**c. The dissimilarity between the brain and behavior modules.** The individual dissimilarity values between the variables can be averaged into the predefined behavior domains (Shafto et al., 2014) and brain networks (Yeo, et al., 2011) to simplify their interpretation. This panel shows the averaged dissimilarity within the modules for $S_1$ (*left*), $S_2$ (*right*).

**d. The dissimilarity of the brain-behavior interaction.** The off-diagonal modules that represent the brain-behavior interaction are emphasized for $S_1$ (*left*), $S_2$ (*right*). This is the novel information contributed by running the CCA analysis between the brain and behavior datasets.

**e. Chord plots for visualizing the flow of contribution between brain-behavior domains.** The chord plots represent the data displayed in **d** after thresholding to show the strongest 25th percentile and squaring the values. The sides of the plot represent the overall contribution to the strongest domain relationships. The bands are scaled so that the larger the colored bands the more similar the domains are in their contributions to the CCA. The left panel represents $S_1$ and the right panel represents $S_2$. The difference between $S_1$ and $S_2$ ($S_d$) is reported in Figure 5c.

The dissimilarity values for brain network nodes and behavioral variables (**Supplemental Figure 5c**) were averaged within the 10 brain networks of $Y_{2011}$ and 7 behavioral domains (**Supplemental Figure 5d**). This procedure identifies three portions of $S_1$. The brain-brain dissimilarity, the behavior-behavior dissimilarities and the brain-behavior dissimilarities. The final analysis was focussed on describing the pattern of results in the brain-behavior interactions (**Supplemental Figure 5e**)

The results for the brain-behavioral dissimilarity matrix were also visualized using a modified chord plot (**Supplementary Figure 5e**; see also **Methods**). The plot allowed us to show how multiple associations between brain and behavior load onto $M_1$

simultaneously (**Supplementary Figure 5f**). Two aspects of the plot should be noted: (A) The individual associations between each functional network and behavioral domains are described by the chords. (B) The number of domains (networks) that each network (domain) contributes to is described by the size of the peripheral segments for each network and domain. The more chords, the more contributions of a network (domain) to the various domains (networks). In other words, the larger the segments of a network (domain) the stronger its overall multivariate association.

As an illustrative example, it is convenient to describe the results in $S_1$ by taking the perspective of the behavioral domains and look at how each domain was associated with the various brain network domains. Results show that in the behavioral domains, memory, language, emotion, and attention dominated the association with the brain networks over clinical scores, motor, and social variables (compare the chord edge size in **Supplementary Figure 5f**). In the brain network domains, the frontoparietal, amygdala, hippocampus, subcortical, and dorsal attention domains dominated $S_1$.

Next, the influence of subjects' age was evaluated more specifically by computing the RSA using $M_2$. A dissimilarity matrix $S_2$ was computed repeating the procedure explained above and then $S_1$ and $S_2$ compared.

| Behavior | Strongest Associated Brain Network Modules (Strongest -> Weakest) | | | | | | | | | |
|---|---|---|---|---|---|---|---|---|---|---|
| **Attention** | Hippocampus | Amygdala | Frontoparietal | Subcortical | Dorsal Attention | Somatomotor | Visual | Ventral Attention | DMN | Limbic |
| **Clinical Scores** | Hippocampus | Frontoparietal | Amygdala | Subcortical | Dorsal Attention | Visual | Somatomotor | DMN | Ventral Attention | Limbic |
| **Emotion** | Hippocampus | Frontoparietal | Amygdala | Subcortical | Dorsal Attention | Visual | Somatomotor | DMN | Ventral Attention | Limbic |
| **Language** | Hippocampus | Frontoparietal | Amygdala | Subcortical | Dorsal Attention | Visual | Somatomotor | Ventral Attention | DMN | Limbic |
| **Memory** | Hippocampus | Subcortical | Frontoparietal | Amygdala | Dorsal Attention | Visual | Somoatomotor | Ventral Attention | DMN | Limbic |
| **Motor** | Hippocampus | Frontoparietal | Amygdala | Subcortical | Dorsal Attention | Somatomotor | Visual | DMN | Ventral Attention | Limbic |
| **Social** | Dorsal Attention | Frontoparietal | Hippocampus | Visual | Amygdala | Subcortical | Somatomotor | DMN | Ventral Attention | Limbic |

**Supplemental Table 1.** The ranked order of the dissimilarity between behavior domains to network modules.
